# Enhancer-promoter interactions form independently of genomic distance and are functional across TAD boundaries

**DOI:** 10.1101/2022.08.29.505755

**Authors:** Pedro Borges Pinto, Alexia Grasso, Deevitha Balasubramanian, Séverine Vincent, Hélène Tarayre, Damien Lajoignie, Yad Ghavi-Helm

## Abstract

Developmental enhancers are essential regulatory elements that drive precise spatio-temporal gene expression patterns. They do so by interacting with the promoter of their target genes, often across large genomic distances, in a highly specific manner. However, it is unclear how such specificity can be achieved. While several studies have suggested that Topologically Associating Domains (TADs)^1–3^ facilitate and constrain enhancer-promoter interactions, the role of TAD boundaries in effectively restricting enhancer-promoter interactions is heavily debated. Here we show that enhancers can establish long-range interactions across TAD boundaries and even between different chromosomes. Moreover, some of these interactions are functional *in vivo*, illustrating their functional importance. Using the *twist* locus in *Drosophila* embryos, we systematically relocated one of its enhancers to different regulatory contexts and distances from the *twist* promoter. We found that the *twist* promoter can engage in functional enhancer-promoter interactions across a TAD boundary and that distal interactions are sometimes favored over proximal ones. Our results demonstrate that TAD boundaries are not sufficient to constrain enhancer-promoter interactions and that the formation of long-range interactions is not solely driven by distance. These observations suggest that other general mechanisms must exist to establish and maintain specific enhancer-promoter interactions across large distances.

---

Enhancers are short non-coding genomic elements that play a crucial role in the regulation of gene expression during development, by driving precise spatial and temporal expression patterns^4^. They can be located at various distances from the promoter of their target gene(s), sometimes even skipping multiple nearby promoters to regulate the expression of a gene located at a large genomic distance^5,6^. Long-range enhancer-promoter interactions are mediated through the formation of three-dimensional (3D) chromatin loops^7^. Even in the compact *Drosophila* genome, the distance between enhancers and their target genes is comparable to mammals, with a median distance of 100 kb, and some interactions spanning distances of over 500 kb^8^. In this context, it is essential for enhancers to target and regulate the expression of the correct gene while avoiding the inappropriate expression of neighboring genes.

In recent years, genome topology has been suggested to play an important role in constraining enhancer-promoter interactions. Indeed, these interactions tend to be constrained within large regulatory domains which broadly coincide with regions of increased three-dimensional proximity named Topologically Associating Domains (TADs)^9–13^. Rearrangements affecting TAD boundaries can impair proper enhancer-promoter communication and affect gene expression^14–21^. TADs have thus been proposed to act as functional regulatory units that favor local enhancer-promoter interactions whilst preventing interactions across their boundaries^22,23^. However, in some cases, gene expression seems resilient to chromosomal rearrangements^24–27^. Moreover, depleting the complexes responsible for TAD boundary formation completely abolishes TAD structures, yet only mildly affects gene expression^28–31^. These conflicting observations question the role of TADs in gene expression regulation^32^. In particular, to what extent can enhancers interact and regulate the expression of their target gene across TAD boundaries, irrespective of genomic distance and chromatin context, and how the specificity of such long-range interactions is achieved remain open questions.

To address these questions, we searched for a well-characterized developmental gene whose activity is easily tractable during embryogenesis and whose expression is regulated by a tissue-specific enhancer. We, therefore, focused on the *Drosophila twist* (*twi*) gene, that codes for a highly conserved transcription factor acting as a master regulator of mesoderm development and promoting epithelial-mesenchymal transition in normal and metastatic cells^33^. In the embryo, *twist* is strongly expressed from the onset of zygotic transcription in the ventral region of the embryo corresponding to the mesoderm anlage and starts to decline after germ band elongation^34^. *twist* mutants are recessive lethal due to abnormal gastrulation characterized by the absence of mesoderm derivatives^35^. During early embryogenesis, the expression of the *twist* gene is regulated by three known enhancers: an upstream distal enhancer (DE), an upstream proximal enhancer (PE)^36^, and a downstream distal enhancer^37^. For simplicity, we will hereafter refer to these enhancers as E1, E2, and E3, respectively (Fig. 1a). Previous reports suggested that these regulatory regions might be active in overlapping cell types during the early stages of embryogenesis^38–40^. Our detailed analysis of the activity of these enhancers revealed that all three regulatory regions are active until stage 10. However, at stage 11, we discovered that E3 is the only active regulatory region controlling *twist* expression in the thoracic and abdominal region (Fig. 1b and Extended Data Fig. 1b). Deleting the endogenous E3 enhancer causes a recessive lethal phenotype at embryonic stages. It is associated to a strong reduction in *twist* expression at stage 11 leading to severe defects in the embryonic somatic musculature of *twi*^*ΔE3*^ mutant embryos (Fig. 1c-d, Extended Data Fig. 1c-d). We, therefore, concluded that the E3 enhancer is essential for the proper development of the mesoderm and that its activity is not redundant with that of the upstream E1 and E2 enhancers.

**Fig. 1:**
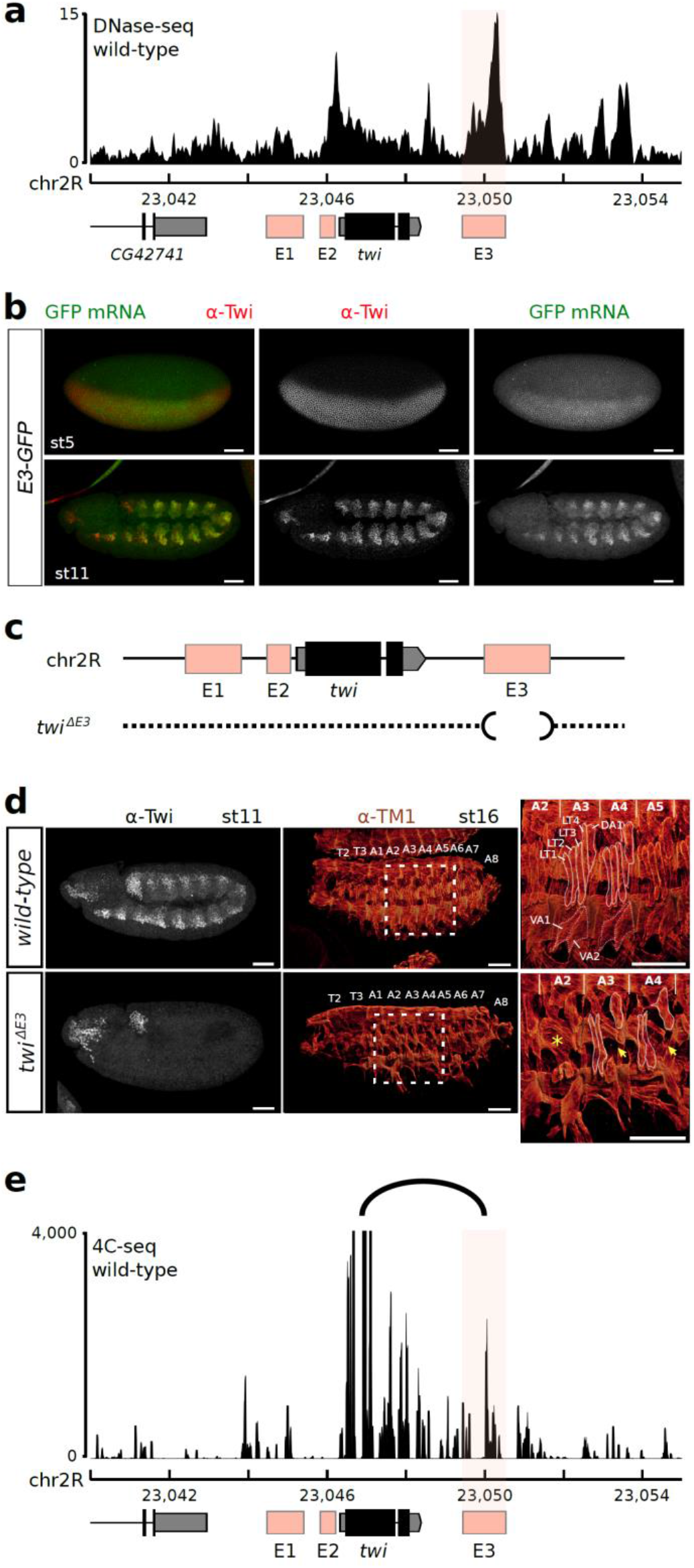
The E3 enhancer activates the expression of *twist* during embryogenesis. **a**. The twist E3 enhancer is located in an open chromatin region as defined by DNase-seq signal in wild-type embryos at stage 11^42^. **b**. Immunostaining with the α-Twist antibody (red) and expression (smiFISH) driven by its E3 enhancer (*GFP*, green) at stage 5 (top) and 11 (bottom). Scale bars 50 μm. **c**. Schematic representation of the *twist* locus and of the *twi*^*ΔE3*^ deletion. **d**. Immunostaining with the α-Twist antibody at stage 11 (white, left) and the α-TM1 antibody at stage 16 (red, middle) in wild-type (top) and *twi*^*ΔE3*^ embryos (bottom). The location of thoracic segments T2-T3 and abdominal segments A1-A8 is indicated. A blow-up (right) indicates the location of specific embryonic body muscles (LT1-4: lateral transverse muscles, DA1: dorsal acute, VA: ventral acute) and the location of missing muscles in the mutant (yellow asterisk: muscles absent in the whole segment, yellow arrows: absence of specific muscles). Scale bars 50 μm. **e**. 4C-seq interaction map at the *twist* locus in wild-type embryos at 5 to 8 hours after egg-lay. The observed interaction between the *twist* promoter and the E3 enhancer is highlighted by an arc. One representative experiment is shown.

We next verified that the endogenous E3 enhancer interacts with the *twist* promoter. The *twist* gene and its three enhancers are located within the same TAD, close to its boundary and in an open chromatin region marked by active histone modifications (Extended Data Fig. 1a, Extended Data Fig. 2a). The E3 enhancer is located approximately 3 kb downstream of the *twist* promoter (Extended Data Fig. 1a). To visualize chromatin interactions at such short distances, we significantly improved our 4C-Seq (circular chromosome conformation capture) protocol and applied it to wild-type *Drosophila* embryos at 5 to 8 hours after egg-lay (stage 10-11) using a viewpoint anchored within the *twist* gene. We observed an interaction between the *twist* promoter and a region overlapping the E3 enhancer, confirming that the endogenous E3 enhancer interacts with the *twist* promoter during early embryogenesis (Fig. 1e and Extended Data Fig. 1a).

Having identified a suitable model to measure the effect of an enhancers’ relocation on gene expression and chromatin organization, we performed extensive genomic engineering of the endogenous *twist* locus by inserting the E3 enhancer at various linear distances from the *twist* promoter (ranging from 7.5 kb to 1.6 Mb and on another chromosome; Fig. 2a). In all cases, the obtained fly lines were homozygous viable. The insertion sites were selected based on their distance to the endogenous *twist* promoter and location with respect to TADs, chromatin domains, and A/B compartments (Extended Data Fig. 2-3). To minimize deleterious effects, we avoided regions containing annotated genes and regulatory sequences (annotated in the REDfly database^41^ or overlapping DNase I hypersensitive sites during embryogenesis^42^) (Extended Data Fig. 2-3). In addition, we ensured that none of the insertion sites interact with the *twist* promoter in wild-type embryos as visualized by 4C-seq (Extended Data Fig. 4 a-b) and Micro-C (Extended Data Fig. 2-3). The E3(+7.5kb) insertion site is located in the same TAD as *twist*, while the E3(+39kb) and E3(+51kb) insertion sites are both located in the adjacent downstream TAD. Two additional insertion sites, E3(−181kb) and E3 (−1.6Mb), are located in more distal upstream regions, and the last insertion site, E3(chr3L), is located on a different chromosome (Fig. 2a, Extended Data Fig. 2-3). Finally, we verified the location of each insertion site with regard to A/B compartments, respectively associated to open and closed chromatin, and established by calculating the eigenvector of a Hi-C contact matrix obtained from stage 5 to 8 whole embryos^43^ (Methods). Except for E3(+51kb), all insertion sites are located in an A compartment.

**Fig. 2:**
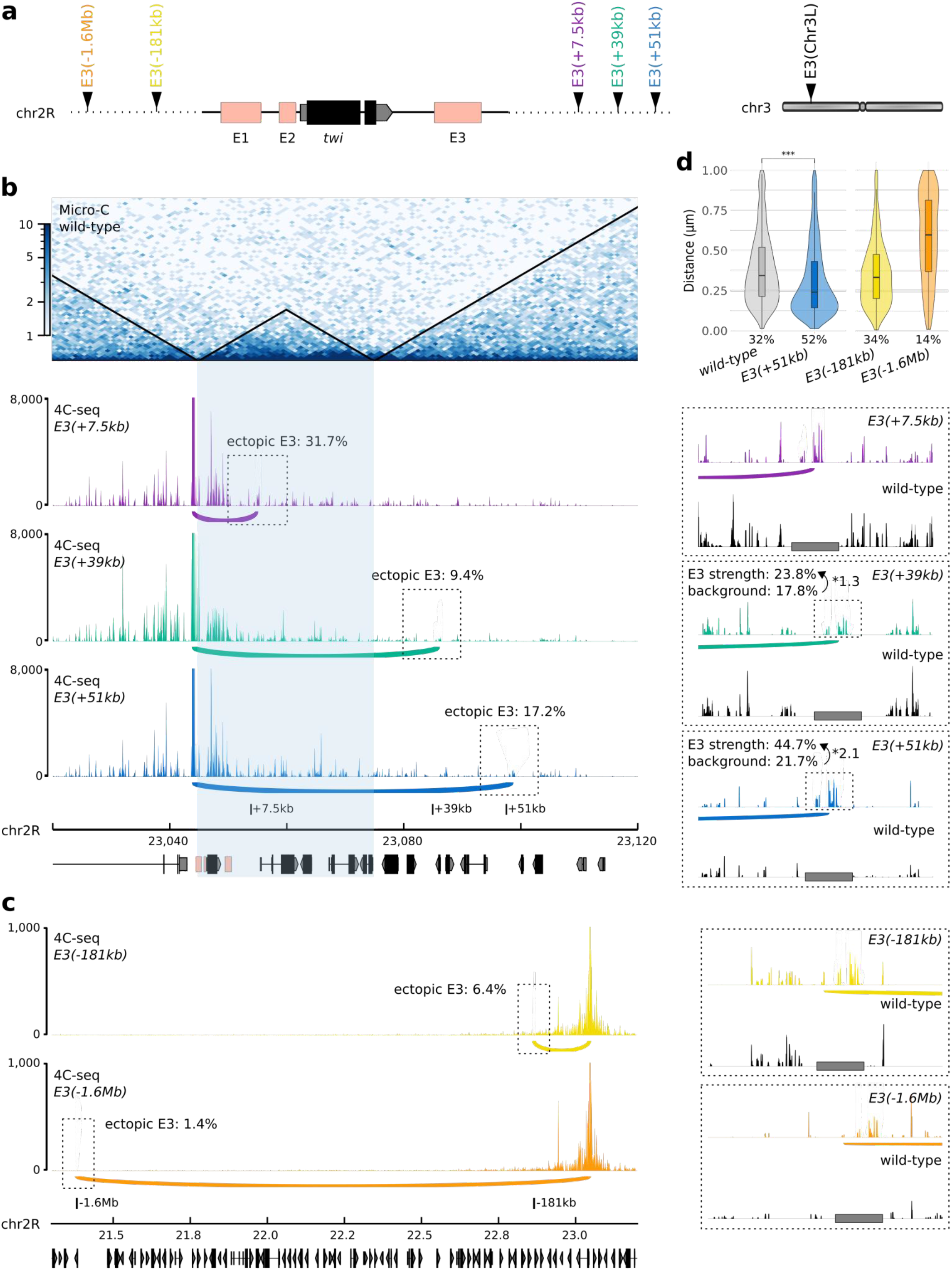
The *twist* promoter interacts with the E3 enhancer across large genomic distances. **a**. Schematic representation of the different ectopic E3 insertion sites on chr2R and chr3L. **b**. High-resolution chromatin organization around the *twist* locus. Top to bottom: normalized Micro-C contact map at 1000 bp resolution in wild-type embryos at 5 to 8 hours after egg-lay (two biological replicates merged), 4C-seq interaction maps in *E3(+7*.*5kb)* (purple), *E3(+39kb)* (green), and *E3(+51kb)* (blue) embryos at 5 to 8 hours after egg-lay (one representative experiment is shown). The TAD containing *twist* and the E3 enhancer is highlighted in light blue. A 10 kb region surrounding the ectopic E3 sites is highlighted by a dotted box and shown as an inset (right). Potential interactions between the *twist* promoter and the ectopic E3 enhancer are highlighted by an arc. The percentage of ectopic E3 reads (ectopic E3), the percentage of reads mapping on the 2-kb ectopic E3 over a 10-kb window (E3 strength), and the percentage of reads mapping on an adjacent control region (background) are indicated. Insets: 4C-seq interaction maps in a 10 kb region around the ectopic E3 sites in *E3(+7*.*5kb)* (purple), *E3(+39kb)* (green), and *E3(+51kb)* (blue) embryos compared to wild-type (black) embryos at the same stage. The location of the E3 ectopic insertion is indicated by a grey rectangle. **c**. 4C-seq interaction maps in *E3(−181kb)* (yellow) and *E3(−1*.*6Mb)* (orange) embryos at 5 to 8 hours after egg-lay (one representative experiment is shown). A 10 kb region surrounding the ectopic E3 sites is highlighted by a dotted box and shown as an inset (right). Potential interactions between the *twist* promoter and the ectopic E3 enhancer are highlighted by an arc. The percentage of ectopic E3 reads (ectopic E3) is indicated. Insets: 4C-seq interaction maps in a 10 kb region around the ectopic E3 sites in *E3(−181kb)* (yellow) and *E3(−1*.*6Mb)* (orange) embryos compared to wild-type (black) embryos at the same stage. The location of the E3 ectopic insertion is indicated by a grey rectangle. **d**. Left: Violin plots representing 3D DNA FISH distances measured in mesodermal nuclei between a probe located next to the *twist* promoter and a probe located next to the +51kb insert site in wild-type (grey) and *E3(+51kb)* (blue) embryos at stage 11. A non-parametric two-sample Kolmogorov–Smirnov test was used to assess the significant difference between DNA FISH distance distributions (p = 2.2e^-16^). Right: Violin plots representing 3D DNA FISH distances measured in mesodermal and non-mesodermal nuclei between a probe located next to the *twist* promoter and a probe located next to the -181kb insert site in *E3(−181kb)* (yellow) embryos and to the -1.6Mb insert site in *E3(−1*.*6Mb)* (orange) embryos at various *twist*-expressing stages. The percentage of colocalization (defined as the percentage of probe pairs with a distance < 0.25 μm; Methods) is indicated for each condition.

We initially used these six fly lines to establish whether ectopically inserted E3 enhancers could engage in long-range enhancer-promoter interactions with the *twist* promoter (Fig. 2b-d). For this purpose, we conducted a comprehensive analysis of chromatin organization in these fly lines, by generating 4C-seq interaction maps from *Drosophila* embryos collected at 2 to 5 hours (stage 5-9) and 5 to 8 hours (stage 10-11) after egg-lay. To have a more comprehensive view of chromatin organization, we used two different viewpoints: one located at the TAD boundary upstream of the *twist* promoter (viewpoint Twi1) and one within the *twist* gene (viewpoint Twi2). In these lines, the endogenous and ectopic copies of the E3 enhancer contain several naturally-occurring single-nucleotide variants (Extended Data Fig. 5) enabling us to differentiate endogenous and ectopic interactions in genomic experiments.

We measured three parameters in our 4C-seq maps, to characterise the interactions of the *twist* locus with each ectopic E3 insertion: i) “ectopic E3”, defined as the percentage of interactions exclusively established with the ectopic version of the E3 enhancer over all versions of E3. ii) “E3 strength”, defined as the percentage of interactions overlapping the ectopic enhancer site as compared to a 10 kb region around the insertion site and used to estimate the strength of the interaction. iii) “background”, defined as the percentage of interactions overlapping a sliding window around (but excluding) the ectopic enhancer site as compared to a 10 kb region around the insertion site and used to estimate the expected background level of interactions at this site. We considered two regions as interacting if ectopic E3 was greater than 10% and/or E3 strength/background greater than 1.7.

We first focused on the fly lines carrying an insertion site downstream of the *twist* locus, corresponding to the insertions at +7.5 kb (line *E3(+7*.*5kb)*), +39 kb (line *E3(+39kb)*), and +51 kb (line *E3(+51kb)*) from the *twist* promoter. Placing the ectopic E3 enhancer in the same TAD as *twist* (line *E3(+7*.*5kb)*) did not affect its ability to engage in chromatin interactions with the *twist* promoter. Indeed, on average 31.7% of the reads mapping to the E3 enhancer at 5-8 h after egg-lay corresponded to the ectopic E3 enhancer, indicating that the *twist* promoter interacts with both copies of the E3 enhancer (Fig. 2b, Extended Data Fig. 6a, c).

When the ectopic E3 enhancer was inserted in a different TAD, however, we observed two opposite situations. When inserted at position +39kb (line *E3(+39kb)*), only 9.36% of the reads mapped to the ectopic E3 enhancer at 5-8 h after egg-lay, and no significant interaction was observed when compared to a wild-type control (Fig. 2b, Extended Data Fig. 6b-d). In contrast, when the ectopic E3 enhancer was inserted at position +51kb (line *E3(+51kb)*), 17.2% of the reads map to the ectopic E3 enhancer (Fig. 2b, Extended Data Fig. 6c, Extended Data Fig. 7a). In addition, the interaction strength (“E3 strength”) was nearly doubled in the *E3(+51kb)* line (44.7%) compared to the *E3(+39kb)* line (23.8%) or their respective background controls (21.7% and 17.8% respectively) (Fig. 2b, Extended Data Fig. 6d). We also observed an increase in the ectopic E3 interaction frequency between 2 to 5 and 5 to 8 hours after egg-lay, from 26% to 31.7% in line *E3(+7*.*5kb)* and from 13.7% to 17.2% in line *E3(+51kb)*. This was not the case for the non-interacting *E3(+39kb)* line (from 9.8% to 9.4%; Extended Data Fig. 6c). This increase correlates with the specific activity of E3 during embryogenesis at stage 11. We further validated the interaction between the *twist* promoter and the ectopic E3(+51kb) enhancer using mesoderm-specific 3D DNA fluorescence *in situ* hybridization (FISH) in *E3(+51kb)* embryos at stage 5 and 11 by measuring the distance between a probe located near the *twist* promoter and the ectopic E3 insertion. As a control, we measured the same distance in a wild-type line. The distance distribution was significantly different between the two lines (p = 2.2e^-16^), with a percentage of colocalization increasing from 32% in the non-interacting control to 52% in the *E3(+51kb)* line (Fig. 2d, Extended Data Fig. 8a-b).

In the lines *E3(−181 kb)* and *E3(−1*.*6Mb)*, the ectopic E3 enhancer is located at a much larger distance from the *twist* locus, with several TAD boundaries in between. In these lines, we observed that on average less than 10% of the reads mapping to the E3 enhancer corresponded to the ectopic E3 (Fig. 2c). Besides, the ectopic E3 site was not enriched in 4C-seq interactions in those fly lines compared to a wild-type control (Fig. 2b-c - inset). This absence of interaction was also validated by 3D DNA FISH in the *E3(−1*.*6Mb)* and *E3(−181kb)* fly lines (Fig. 2d; 34% and 14% colocalization, respectively), confirming that the *twist* promoter is not able to engage in long-range enhancer-promoter interactions with the E3 enhancer when it is inserted at the -1.6Mb and -181kb sites. Together, our observations suggest that the *twist* promoter can engage in long-range enhancer-promoter interactions with the E3 enhancer in a distance-independent manner, with distal sites (for eg. the +51kb site) sometimes favored over more proximal ones (for eg. the +39kb site).

To demonstrate the biological relevance of the observed interactions between the *twist* promoter and various ectopic E3 insertions, we probed to what extent the ectopic insertions could rescue the deletion of the endogenous E3 enhancer (*twi*^*ΔE3*^) (Fig. 3a-b, Extended Data Fig. 9a). In line with our previous observations, inserting the ectopic E3 enhancer 7.5kb downstream of the *twist* promoter, fully rescued the viability (Fig. 3b, *twi*^*ΔE3*^, *E3(+7*.*5kb)*, Extended Data Fig. 9a; 81% versus 0%), *twist* expression (Extended Data Fig. 9b), and muscle formation (data not shown) of *twi*^*ΔE3*^ embryos. There was however a more modest effect upon the insertion of the ectopic E3 enhancer 51 kb away from the *twist* promoter, with *twi*^*ΔE3*^, *E3(+51kb)* embryos displaying a partial rescue of embryo viability (Fig. 3b, Extended Data Fig. 9a; 14% versus 0%), *twist* expression (Fig. 3c, Extended Data Fig. 9b), and muscle formation (Fig. 3d) relative to *twi*^*ΔE3*^ embryos. The rescue of *twist* expression in *twi*^*ΔE3*^, *E3(+51kb)* embryos followed two different patterns (Extended Data Fig. 9b): in 29% of the embryos, *twist* was expressed at low levels throughout the whole mesoderm. In the other 61% of the embryos, however, *twist* was expressed at a higher level, but was restricted to a group of cells in the T1 and maxillary segments of the embryo. Finally, about 10% of the embryos displayed both a weak *twist* expression throughout the mesoderm and a high expression in the group of cells. In contrast, none of the non-interacting insertions were able to rescue embryo viability (Extended Data Fig. 9a). These results further support the functionality of the long-range enhancer-promoter interactions we identified.

**Fig. 3:**
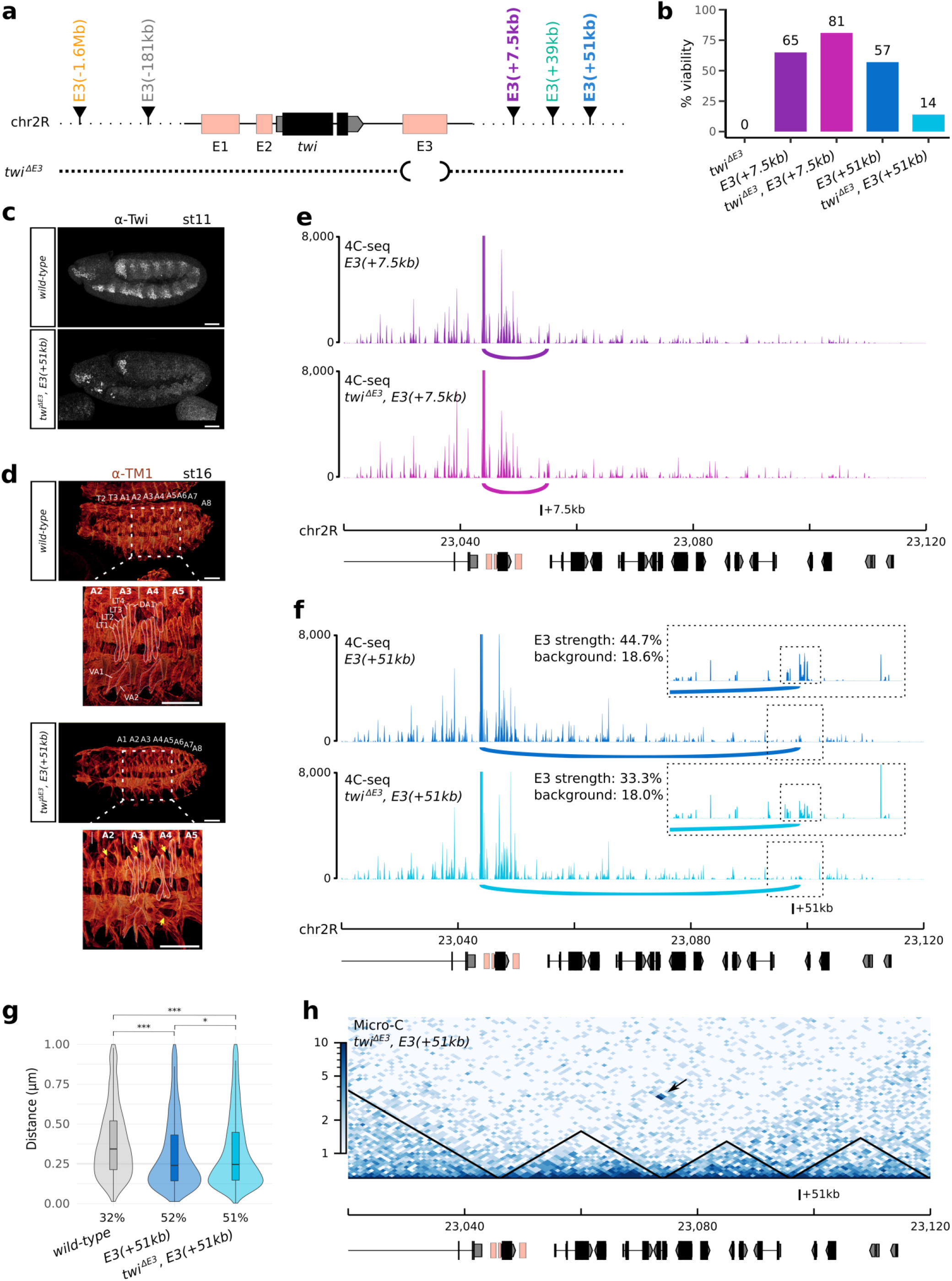
Long-range interactions between the ectopic E3 and the *twist* promoter can rescue *twi*^*ΔE3*^ mutants. **a**. Schematic representation of the different ectopic E3 insertion sites on chr2R and of the *twi*^*ΔE3*^ deletion. **b**. Bar plot representing the percentage of viable embryos on the *twi*^*ΔE3*^ (black), *E3(+7*.*5kb)* (dark purple), *twi*^*ΔE3*^, *E3(+7*.*5kb)* (light purple), *E3(+51kb)* (dark blue), and *twi*^*ΔE3*^, *E3(+51kb)* (light blue) lines. For each condition, at least two independent experiments were performed, with at least 50 embryos each. **c**. Immunostaining with the α-Twist antibody in wild-type (top) and *twi*^*ΔE3*^, *E3(+51kb)* embryos (bottom) at stage 11. Scale bars 50 μm. **d**. Immunostaining with the α-TM1 antibody in wild-type (top) and *twi*^*ΔE3*^, *E3(+51kb)* embryos (bottom) at stage 16. The location of thoracic segments T2-T3 and abdominal segments A1-A8 is indicated. A blow-up (dotted scare) indicates the location of specific embryonic body muscles (LT1-4: lateral transverse muscles, DA1: dorsal acute, VA: ventral acute) and the location of missing muscles in the mutant (yellow arrows). Scale bars 50 μm. **e**. 4C-seq interaction maps in *E3(+7*.*5kb)* (dark purple) and *twi*^*ΔE3*^, *E3(+7*.*5kb)* (light purple) embryos at 5 to 8 hours after egg-lay (one representative experiment is shown). **f**. 4C-seq interaction maps in *E3(+51kb)* (dark blue), and *twi*^*ΔE3*^, *E3(+51kb)* (light blue) embryos at 5 to 8 hours after egg-lay (one representative experiment is shown). A 10 kb region surrounding the ectopic E3 sites is highlighted by a dotted box and shown as an inset. Potential interactions between the *twist* promoter and the ectopic E3 enhancer are highlighted by an arc. The percentage of reads mapping on the 2-kb ectopic E3 over a 10-kb window (E3/bkgd) and the percentage of reads mapping on an adjacent control region (control) are indicated. **g**. Violin plots representing 3D DNA FISH distances measured in mesodermal nuclei between a probe located next to the *twist* promoter and a probe located next to the +51kb insert site in wild-type (grey), *E3(+51kb)* (dark blue), and *twi*^*ΔE3*^, *E3(+51kb)* (light blue) embryos at stage 11. A non-parametric two-sample Kolmogorov–Smirnov test was used to assess the significant difference between DNA FISH distance distributions (wild-type versus *E3(+51kb):* p = 2.2e^-16^, wild-type versus *twi*^*ΔE3*^, *E3(+51kb):* p = 2.2e^-16^, *E3(+51kb)* versus *twi*^*ΔE3*^, *E3(+51kb):* p = 0.04). The percentage of colocalization (defined as the percentage of probe pairs with a distance < 0.25 μm; Methods) is indicated for each condition. **h**. Normalized Micro-C contact map at 1000 bp resolution in *twi*^*ΔE3*^, *E3(+51kb)* embryos at 5 to 8 hours after egg-lay (two biological replicates merged). The interaction between the *twist* locus and the +51kb insertion site is indicated by a black arrow.

To confirm that the activation of *twist* by the ectopic E3 enhancers is dependent on enhancer-promoter looping, we repeated 4C-seq experiments in *twi*^*ΔE3*^, *E3(+7*.*5kb)* and *twi*^*ΔE3*^, *E3(+51kb)* embryos, where the endogenous E3 was deleted. In both cases, the interaction between the *twist* promoter and the +7.5 kb or +51 kb ectopic E3 enhancers was maintained (Fig. 3 e-f, Extended Data Fig. 10a-b). Mesoderm-specific 3D DNA FISH experiments performed at stage 5 and 11 further validated this observation (Fig. 3g, Extended Data Fig. 8a), with a significant difference in the distance distribution (p = 2.2e^-16^) and an increased colocalization from 32% in the wild-type control to 51% in the *twi*^*ΔE3*^, *E3(+51kb)* line. Deleting the endogenous E3 enhancer in the *E3(+51kb)* line however resulted in a slight decrease in 4C-seq interaction frequency (Fig. 3f; E3 strength from 44.7% to 33.3%) as well as a slight increase in the distance between the *twist* promoter and the +51kb insertion site (Extended Data Fig. 8a; colocalization from 56% to 43%). This effect appeared more pronounced at stage 5 than at stage 11 (Extended Data Fig. 8a), suggesting that enhancer-enhancer interactions between the two copies of E3 might favor a more compact conformation at earlier stages. Together, these data show that the *twist* promoter can engage in functional enhancer-promoter interactions across large distances (*i*.*e*. over distances greater than about 10 kb, which would be the typical average distance between known enhancer-promoter pairs in *Drosophila*) and that these interactions do not depend on the presence of the endogenous E3 enhancer.

To establish whether long-range enhancer-promoter interactions indeed can take place across TAD boundaries and analyse global changes in chromatin organisation, we generated Micro-C contact maps at 5 to 8 h after egg-lay in the wild-type and *twi*^*ΔE3*^, *E3(+51kb)* lines. The Micro-C maps provided us with an additional opportunity to confirm the presence of long-range interactions between the *twist* locus and the ectopic E3 enhancer at the +51kb site (Fig3h, black arrow) that were absent in the wild-type line (Fig. 2b, Extended Data Fig. 11). Ectopically inserting the E3 enhancer at position +51kb leaded to increased local interactions between these two sites, yet did not significantly alter global chromatin organization (Fig3h, Extended Data Fig. 11). Indeed, this ectopic insertion did not affect the presence of the boundary located between the *twist* locus and the +51kb site, but instead drove the formation of an additional boundary at the +51kb site, confirmed by a strong shift in the directionality index (Extended Data Fig. 11, blue arrow). This insertion also resulted in a shift of the region between the *twist* locus and the +51kb site from an A compartment to a B compartment, while the position of the +51kb site itself shifted from a B compartment to an A compartment (Extended Data Fig. 11, red arrow). These results confirm that a long-range functional interaction is established between the *twist* promoter and the ectopic E3 enhancer at position +51kb and that this interaction takes place across a TAD boundary. This long-range cross-TAD regulation challenges the current view of enhancer biology, whereby enhancer-promoter interactions are constrained by TAD boundaries.

The last E3 insertion site included in our analysis is located on a different chromosome than *twist*, allowing us to investigate the presence of inter-chromosomal enhancer-promoter interactions. Surprisingly, 4C-seq experiments in the *E3(chr3L)* line revealed that an average of 46.9% of the reads mapping to the E3 enhancer at 5-8 h after egg-lay corresponded to the ectopic E3 enhancer (Fig. 4a-b, Extended Data Fig. 6c, Extended Data Fig. 12 a-b). The interaction between the *twist* promoter and the ectopic E3(chr3L) enhancer was also validated by mesoderm-specific 3D DNA FISH in *E3(chr3L)* embryos at stage 5 by measuring the distance between a probe located near the *twist* promoter and the ectopic E3 insertion. As a control, we measured the same distance in a wild-type line. The distance distribution was significantly different between the two lines (p = 9.6e^-6^), with a percentage of colocalization increasing from 9% in the non-interacting control to 21% in the *E3(chr3L)* line (Fig. 4c). We observed two populations of nuclei in *E3(chr3L)* embryos: a population where the two loci are very distant (as in the wild-type condition), and a population where the two loci are highly colocalized. This might indicate that, while this interaction can be very strong in some cells, it is also highly unstable. This observation is further supported by the inability of the E3(chr3L) insertion to rescue both the viability (Extended Data Fig. 9a) and *twist* expression in *twi*^*ΔE3*^ mutant embryos (Fig. 4d). Overall, these data indicate that while strong, the ectopic E3(chr3L) enhancer fails to establish functional interactions with the *twist* promoter.

**Figure 4:**
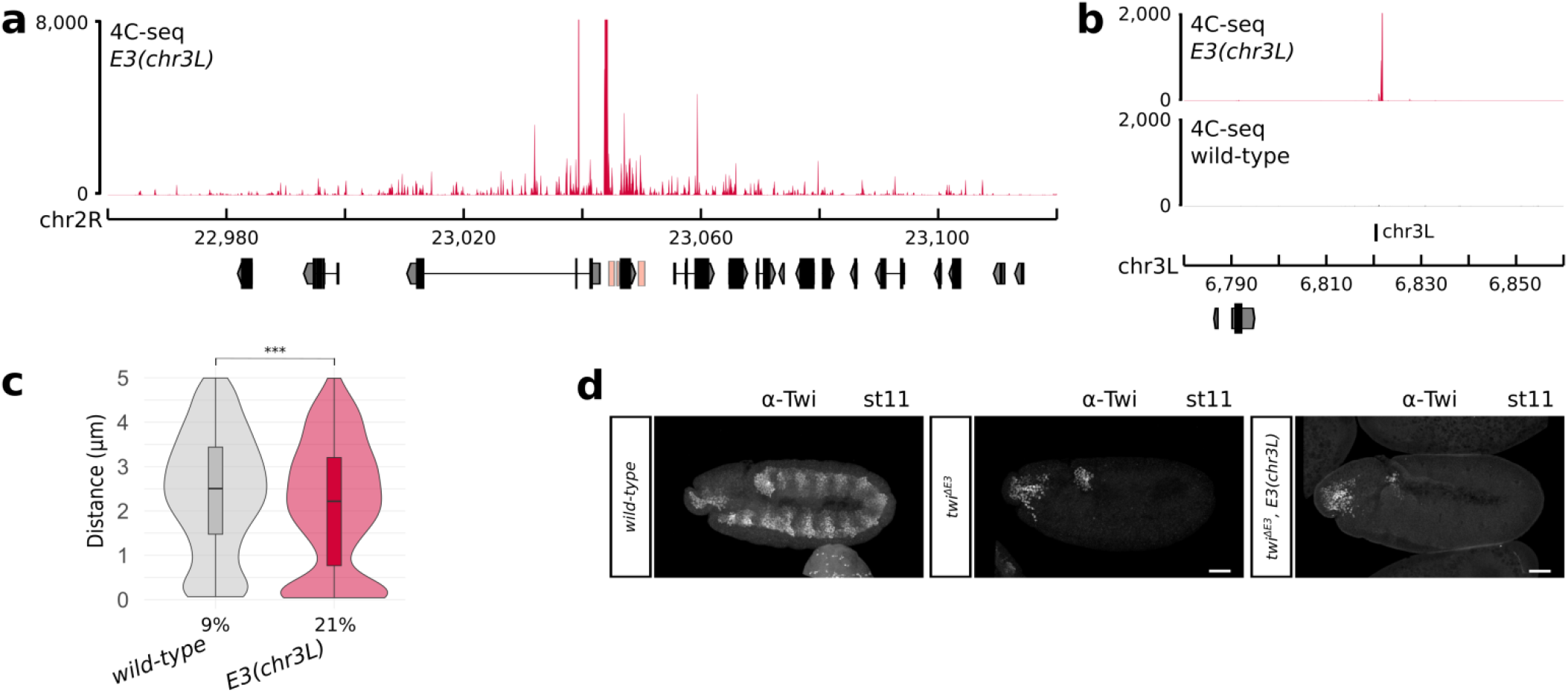
The *twist* promoter interacts with the E3 enhancer placed on a different chromosome. **a**. 4C-seq interaction map in *E3(chr3L)* (red) embryos at 5 to 8 hours after egg-lay around the *twist* locus (one representative experiment is shown). **b**. 4C-seq interaction maps in *E3(chr3L)* (red) and wild-type (black) embryos at 5 to 8 hours after egg-lay around the chr3L insert site (one representative experiment is shown). **c**. Violin plots representing 3D DNA FISH distances measured in mesodermal nuclei between a probe located next to the *twist* promoter and a probe located next to the chr3L insert site in wild-type (grey) and *E3(chr3L)* (red) embryos at stage 5. A non-parametric two-sample Kolmogorov–Smirnov test was used to assess the significant difference between DNA FISH distance distributions (p = 9.6e^-6^). The percentage of colocalization (defined as the percentage of probe pairs with a distance < 0.25 μm; Methods) is indicated for each condition. **d**. Immunostaining with the α-Twist antibody at stage 11 in wild-type (left), *twi*^*ΔE3*^ (middle), and *twi*^*ΔE3*^, *E3(chr3L)* (right) embryos. Scale bars 50 μm.

In summary, by systematically perturbing a specific locus with controlled genetic tools, we uncovered fundamental features of enhancer-promoter interaction specificity across large distances. We demonstrated that the *twist* promoter can engage in long-range interactions with an ectopic enhancer across large distances and across TAD boundaries, and even between chromosomes. Rescue experiments confirmed that such long-range interactions can sometimes be functional during embryonic development and do not depend on the presence of the endogenous enhancer. Our data thus reveal that enhancer-promoter interactions are not necessarily constrained by TAD boundaries. In fact, the *twist* promoter can interact with the E3 enhancer when it is located ectopically at a distance 17 times greater than the endogenous enhancer. This observation is in agreement with previous reports describing minor transcriptional effects upon the disruption of TADs boundaries^24–30^ and cross-TAD transcriptional regulation at the *xist* locus^44^. We also demonstrate that the formation of long-range enhancer-promoter interactions is not solely dependent on the distance between the enhancer and the promoter, as the E3 enhancer can activate *twist* expression when located 51kb away from the promoter but not when located at a more proximal position (39kb). While TAD-mediated enhancer-promoter proximity certainly favors rapid gene activation^45,46^, especially in cells characterized by very fast cell cycles such as *Drosophila* embryonic cells, other mechanisms must exist to promote interactions across large distances. Overall, neither the genomic distance, the location of TAD boundaries, nor the enhancer sequence can solely dictate enhancer-promoter interaction specificity. Instead, we propose that long-range enhancer-promoter interactions would be favored between specific genomic sites, a feature which is reminiscent of architectural proteins such as insulators^47^ and tethering elements^48,49^. However, the presence of such proteins is not sufficient to drive long-range interactions. Indeed, the +51 kb insertion site does not interact with the *twist* promoter in wild-type conditions, but only upon the insertion of the E3 ectopic enhancer. Therefore, we propose a model where enhancer-promoter interaction specificity across large distances is governed by an interplay between the sequence of the enhancer itself and the sequence of the genomic locus where it is inserted.

## METHODS

### Plasmid construction and transgenic fly generation

All plasmids were constructed using standard cloning methods with New England Biolabs restriction enzymes and T4 DNA ligase (New England Biolabs) or with the NEBuilder HiFi DNA Assembly kit (New England Biolabs). All constructs were verified by sequencing.

Unless specified otherwise, “wild-type” fly lines used in this study refer to the *yw y[1] w[1118]* line (BDSC_6598). All fly lines were raised on standard food at 25°C.

To create ectopic E3 insertion lines, we used two strategies: First, we took advantage of the popular MiMIC (*Minos* Mediated Integration Cassette) system^50^, which consists of a *Minos* transposon carrying a *yellow*^*+*^ dominant body-color marker and a gene-trap cassette flanked by two inverted ΦC31 integrase *attP* sites. This cassette can be efficiently replaced by another cassette containing the DNA sequence of interest flanked by two inverted ΦC31 integrase *attB* sites using RMCE. This insertion event can be conveniently identified by the loss of body pigmentation in adult flies (corresponding to the replacement of the *yellow*^*+*^ marker by the sequence of interest). We used this strategy to generate five different fly lines where the E3 enhancer was inserted at different locations. While such MiMIC fly lines are readily available to create insertions at thousands of sites, we had to use a second strategy to specifically insert the E3 enhancer in the same TAD as *twist* (line *E3(+7*.*5kb)*), a region where no MiMIC fly line is available. In this second strategy, we first used CRISPR-Cas9 mediated HDR to create a fly line where a ΦC31 integrase *attP* site is integrated at the desired location. This site was then used to insert the E3 sequence from a donor vector containing a ΦC31 integrase *attB* site.

#### For the ectopic integration of enhancer E3 +7.5 kb away from the *twist* promoter (to obtain line E3(+7.5kb))

The *pHD-dsRed-attP* vector (Addgene #51019^51^) was used to introduce an *attP* docking site at position +7.5 kb. Homology arms (∼1 kb each) surrounding the insertion site were amplified from genomic DNA of the *w[1118]; PBac{y[+mDint2]=vas-Cas9}VK00027* (BDSC_51324^52^) fly line. The gRNA was designed using the flyCRIPSR target finder^51^ (sequence of the gRNA: GTCGAATGTCGGGCATATCTT) and cloned in the *pU6-BbsI-chiRNA* vector (Addgene #45946^53^) following the flyCRISPR recommendations (https://flycrispr.org/). The vectors were co-injected in-house in embryos of the *w[1118]; PBac{y[+mDint2]=vas-Cas9}VK00027* fly line.

The resulting transgenic line was first crossed to a Cre recombinase-expressing line (BDSC_766) to delete the dsRed marker cassette. The E3 sequence was then inserted at position +7.5 kb (*line E3(+7*.*5kb)*) by co-injecting the p3xP3-EGFP.vas-int.NLS vector (Addgene #60948^54^) and the *pattB* vector (DGRC #1420^55^) containing the E3 sequence (2R:23049440-23050529) amplified from genomic DNA of the *yw y[1] w[1118]* fly line and cloned using the KpnI and XhoI restriction sites.

#### For the ectopic integration of enhancer E3 -1.6 Mb, -181 kb, +39 kb, and +51 kb away from the *twist* promoter and on chromosome 3L (to obtain line *E3(−1*.*6 Mb), E3(−181kb), E3(+39kb), E3(+51kb)*, and *E3(chr3L)*)

The E3 sequence (2R:23,049,440-23,050,529) was amplified from genomic DNA of the *yw y[1] w[1118]* fly line, except for the +51 kb line where the E3 sequence was amplified from the *y[1] w[*]; Mi{y[+mDint2]=MIC}MI01218* (BDSC_55415^50^) fly line.

The PCR product was cloned into the *pBS-KS-attB1-2-PT-SA-SD-0-2xTY1-V5* vector (Addgene #61255^54^) using HindIII and XbaI restriction sites. The resulting vector and p3xP3-EGFP.vas-int.NLS were injected in-house through ΦC31-mediated recombination^56^ in embryos of the following “MiMIC” fly lines^50^:

- *y[1] w[*]; Mi{y[+mDint2]=MIC}MI04814* (BDSC_38170) to obtain line *E3(−1*.*6 Mb)*
- *y[1] w[*]; Mi{y[+mDint2]=MIC}MI01218* (BDSC_55415) to obtain line *E3(−181kb)*
- *y[1] w[*]; Mi{y[+mDint2]=MIC}MI11229* (BDSC_55595) to obtain line *E3(+39kb)*
- *y[1] w[*]; Mi{y[+mDint2]=MIC}MI02100* (BDSC_32829) to obtain line *E3(+51kb)*
- *y[1] w[*]; Mi{y[+mDint2]=MIC}MI10934* (BDSC_55560) to obtain line *E3(chr3L)*

#### For the deletion of the endogenous enhancer E3 in the *w[1118]; PBac{y[+mDint2]=vas-Cas9}VK00027, E3(−1*.*6Mb), E3(+7*.*5kb), E3(+39kb)*, and *E3(+51kb)* fly lines (to obtain line *twi*^*ΔE3*^, *twi*^*ΔE3*^, *E3(−1*.*6Mb), twi*^*ΔE3*^, *E3(+7*.*5kb)*, and *twi*^*ΔE3*^, *E3(+51kb)*)

Two ΦC31 integrase *attP* landing sites were inserted into the *pHD-dsRed* vector (DGRC #1360^51^) using either a BsiWI restriction site or an AgeI and a SpeI restriction site. The resulting vector was used to delete the endogenous E3 sequence (2R:23,049,262-23,050,567). Homology arms (∼1 kb each) surrounding the deletion site were amplified from genomic DNA of the *w[1118]; PBac{y[+mDint2]=vas-Cas9}VK00027* fly line. gRNAs were designed using the flyCRIPSR target finder^51^ and cloned in the *pU6-BbsI-chiRNA* vector (Addgene #45946^53^) following the flyCRISPR recommendations (https://flycrispr.org/). Sequence of the gRNAs:

- GAAATCAAAGACTTGTATAC and GGGGAAAAATATCTTTGCAG for line *w[1118];* *PBac{y[+mDint2]=vas-Cas9}VK00027* and *E3(+7*.*5kb)*
- GAAATCAAAGACTTGTATGC and GGGGAAAAATATCTTTGAAG for line *E3(−1*.*6Mb)* and *E3(+39kb)*
- GAAATCAAAGACTTGTATAC and GGGGGGAAATATCTTTGAAG for line *E3(+51kb)*

The vectors were co-injected in-house (except for *twi*^*ΔE3*^, *E3(+7*.*5kb)* which was generated by the FlyORF Injection Service) in embryos of the following fly lines:

- *w[1118]; PBac{y[+mDint2]=vas-Cas9}VK00027* (to obtain line *twi*^*ΔE3*^)
- *E3(−1*.*6Mb)* to obtain line *twi*^*ΔE3*^, *E3(−1*.*6Mb)*
- *E3(+7*.*5kb)* to obtain line *twi*^*ΔE3*^, *E3(+7*.*5kb)*
- *E3(+39kb)* to obtain line *twi*^*ΔE3*^, *E3(+39kb)*
- *E3(+51kb)* to obtain line *twi*^*ΔE3*^, *E3(+51kb)*

Note: the *E3(−1*.*6Mb), E3(+7*.*5kb), E3(+39kb)*, and *E3(+51kb)* fly lines were first crossed with the *w[1118]; PBac{y[+mDint2]=vas-Cas9}VK00027* line to express Cas9 in the germline.

#### For the deletion of the endogenous enhancer E3 in the *E3(chr3L)* fly line (to obtain line *twi*^*ΔE3*^, *E3(chr3L)*)

The deletion of the endogenous E3 sequence in the line where E3 was ectopically inserted on chromosome 3L was obtained by crossing the *E3(chr3L)* fly line with the *twi*^*ΔE3*^ fly line.

The final coordinates of all insertion sites are as follows:

- line *E3(−1*.*6 Mb)*: 2R:21,381,841
- line *E3(−181kb)*: 2R:22,865,023
- line *E3(+7*.*5kb)*: 2R:23,053,901
- line *E3(+39kb)*: 2R:23,084,945
- line *E3(+51kb)*: 2R:23,097,536
- line *E3(chr3L)*: 3L:6,820,484

#### For transgenic reporter assays

To assess the enhancer activity of the E1, E2, and E3 enhancers, these enhancers were cloned upstream of the minimal *twist* promoter (chr2R:23,046,216-23,046,481) driving a mGFPmut2 reporter gene^57^ (codon-optimized for *Drosophila*) in the pBID vector backbone (Addgene #35190). The coordinates of the cloned regions are as follows: enhancer E1: chr2R: 23,044,478-23,045,403, enhancer E2: chr2R:23,045,827-23,046,215, enhancer E3: chr2R: 23,049,440-23,050,529. All constructs were injected in-house through ΦC31-mediated recombination^56^ into the *nos-ϕC31\int*.*NLS; attP40* line^58^. Stably integrated transgenic lines were balanced, and homozygous lines were used for immunostaining and smiFISH to examine *GFP* expression.

### Embryo collections

Freshly hatched adults of the appropriate genotype were placed in embryo collection vials with standard apple cap plates. *Drosophila* embryos were collected on apple juice agar plates at 25 °C at the appropriate time-point (after 3 pre-lays of 1 hour for stage-specific collections), dechorionated using 50% bleach, and washed alternately with water and PBS + 0.1% Triton X-100. The embryos used for 4C-Seq were covalently crosslinked in 1.8% formaldehyde for 15 min at room temperature and stored at -80°C. The embryos used for Micro-C were covalently crosslinked in 1.8% formaldehyde for 15 min, quenched for 5 min with 2M Tris-HCl pH7.5, then crosslinked again with 3 mM DSG for 45min and stored at -80°C. The embryos used for 3D DNA FISH were covalently crosslinked in 4% formaldehyde for 25 min and stored at -20°C in methanol. The embryos used for immunostaining were covalently crosslinked in 6% formaldehyde for 30 min and stored at -20°C in methanol. The embryos used for smiFISH were covalently crosslinked in 8% formaldehyde for 45 min and stored at -20°C in methanol.

### 4C-seq in *Drosophila* embryos

#### Experimental protocol

Nuclear extraction was carried out as described previously^8^. About 100 to 1000 embryos were used for each 4C template preparation using MboI and NlaIII as the first and second restriction enzymes, respectively. 4C templates were amplified from 320 ng of 4C template using the following primers:

- Twi1_FW: TACGTGCACCAAAAGTTTCTT Twi1_RV: AAAATGGTCGTCAAAGCGC, corresponding to a viewpoint located upstream for the *twist* promoter (chr2R:23,044,043-23,044,500; referred to as viewpoint Twi1)
- Twi2_FW: GGCAACAATCCGAGTGGC Twi2_RV: GTACTCCGAGGGCAGTGG corresponding to a viewpoint located within the *twist* gene (chr2R:23,046,575-23,046,906; referred to as viewpoint Twi2)

An additional 1 to 8 nucleotides “shift” sequence was added at the beginning of the primers, to allow optimal base-pair diversity at the beginning of the read after multiplexing.

The PCR product was purified using SPRIselect beads (Beckman Coulter) and 100 ng of each PCR product was used to generate the final libraries using the NEBNext Ultra II DNA Library Prep Kit for Illumina (New England Biolabs). The libraries were indexed for multiplexing using NEBNext multiplex oligos kit for Illumina (New England Biolabs). A total of 54 libraries were generated, with two independent biological replicates for each sample. The libraries were multiplexed and sequenced on a NextSeq500 sequencer (Illumina) using 75-bp paired-end reads (at the IGFL sequencing facility), yielding a total of at least 10 million reads per sample.

#### Data analysis

The quality of the 4C-seq data was confirmed using the FastQC software (http://www.bioinformatics.babraham.ac.uk/projects/fastqc). Adapter sequences were trimmed using TrimGalore (https://www.bioinformatics.babraham.ac.uk/projects/trim_galore/), and the 5’ shift sequence and the primer sequence was trimmed up to the location of the first restriction site (the restriction enzyme cutting site was kept) using Cutadapt version 2.10^59^. As sequencing was performed in paired-end mode, the fastq files corresponding to each pair were merged into a single fastq file. The trimmed reads were then aligned either to the dm6 reference genome, or to a custom genome, generated using the reform Python tool (https://github.com/gencorefacility/reform). Custom genomes consist of the *Drosophila melanogaster* reference genome (dm6) where the endogenous E3 enhancer sequence has been deleted and re-introduced at the appropriate location, with the appropriate sequence (Extended Data Fig. 5). Six different custom genomes were thus created for each insertion site. Alignment was performed using Bowtie version 1.2.2^60^. As the read length is relatively long, it is possible to obtain reads that result from multiple ligation events between different regions of the genomes. As such reads will not map efficiently to the genome, unmapped reads were retrieved and scanned from the 5’ side for the presence of the first, then of the second restriction site, and trimmed after this location. As a consequence, the reads typically start with the sequence of the first restriction enzyme and have either the sequence of the first or the second restriction site at their 3′ end. These trimmed reads were remapped as previously described and both alignments were merged. The libraries were then normalized by scaling using the read-per-million method and transformed into coverage bedgraph files. 4C-seq data were plotted using pyGenomeTracks version 3.6^61,62^. Reads mapping to the ectopic version of E3 were differentiated from those mapping to the endogenous E3 using single nucleotide variants (highlighted by a red asterisk in Extended Data Fig. 5). This was used to compute the percentage of reads mapping to the ectopic version of E3, out of all reads mapping to some version of E3 (“ectopic E3”). For visualization purposes, the signal displayed over the ectopic E3 regions in 4C interaction maps corresponds to a down-sampling of the overall E3 reads to fit the percentage of ectopic reads. The “strength” of the ectopic E3 signal was estimated by calculating the ratio of reads overlapping the ∼ 2 kb ectopic E3 region over the total number of reads in a 10 kb region around the insertion site (“E3 stregth”). This ratio was compared to a control ratio calculated as the average over six similar windows located slightly upstream or downstream of the ectopic E3 insert site (“background”). Each biological replicate was analyzed independently, and the statistics where averaged across the replicates.

### Micro-C in *Drosophila* embryos

#### Experimental protocol

Micro-C libraries were generated based on a previously established protocol^63^, with appropriate modifications for *Drosophila* embryos.

Nuclear extraction was carried out as described previously^8^ using cold MB#1 buffer (50 mM NaCl, 10 mM Tris-HCl pH 7.5, 5 mM MgCl_2_, 1 mM CaCl_2_, freshly added 0.2% NP-40 and 1x PIC) to resuspend the embryos and extract the nuclei by Dounce homogenization. Nuclei from each replicate were first used for a test Micrococcal Nuclease (MNase, Worthington Biochemical) titration: Nuclei corresponding to 600 ng of chromatin was digested with 45 U of MNase, and then checked for a yield of 300 ng of chromatin with a 90% mononucleosome / 10% dinucleosome ratio^63^. If required, the original 600 ng of chromatin was adjusted to obtain an appropriate yield. Following these tests, for each replicate, 7 parallel reactions of MNase were set up with the appropriate amount of chromatin and taken forward for end-chewing, end-labeling, and proximity ligation^63^. After reverse cross-linking, 150-200 ng of the obtained chromatin was size-selected for ligated mononucleosomes (250-400 bp) on a 3.5% NuSieve agarose gel. The size-selected chromatin was then pulled-down with 25 μl of streptavidin beads, and taken forward for library preparation using the NEBNext Ultra II DNA Library Prep Kit for Illumina (New England Biolabs). A total of 4 libraries were generated, with two independent biological replicates for each sample. The libraries were sequenced on a NovaSeq sequencer (Illumina) using 150 bp paired-end reads, yielding at least 100 million reads per sample.

#### Data analysis

The quality of the Micro-C data was confirmed using the FastQC software (http://www.bioinformatics.babraham.ac.uk/projects/fastqc). Adaptor sequences were trimmed using TrimGalore (https://www.bioinformatics.babraham.ac.uk/projects/trim_galore/). The paired-end files were then aligned to the dm6 reference genome or a custom genome using the Burrows-Wheeler Alignment (BWA) – maximal exact matches (MEM) tool v7.17-4^64^ (http://bio-bwa.sourceforge.net/bwa.shtml#12). The pairtools v0.3.0 (https://github.com/open2c/pairtools) pipeline was used to detect ligation junctions and quality control the paired sequences: pairtools parse was used to detect the ligation events, pairtools sort was used to block sort the reads, pairtools dedup was used to remove pairs that are PCR duplicates of each other. Pairtools split was then used to generate a Pairs file. Genome-wide contact matrices were generated both using pairix (https://github.com/4dn-dcic/pairix) and cooler^65^, and using Juicer^66^. HiCExplorer v2.2.1.1^67^ was used to detect and remove genomic regions with low signal or with high noise, and to implicitly address biases in the data by normalizing the matrices using the Knight-Ruiz balancing algorithm. The resultant matrices were merged and used to detect TAD boundaries and compute the insulation score using HiCExplorer. The insulation score was and the directionality index were also calculated using FAN-C 0.9.23^68^. A/B compartments were detected using the eigenvector command in Juicer^66^.

### Two-colour 3D DNA FISH (fluorescent *in situ* hybridization)

3D DNA FISH was performed as previously described^69^. Five probe sets were designed, mapping to regions of genomic DNA directly adjacent to the *twist* promoter and the different insertion sites:

- chr2R: 23,040,112-23,051,659 for the *twist* promoter
- chr2R: 21,376,545-21,386,746 for the -1.6Mb insertion site
- chr2R: 22,860,007-22,870,484 for the -181 insertion site
- chr2R: 23,093,753-23,101,622 for the +51kb insertion site
- chr3L: 6,816,164-6,824,775 for the chr3L insertion site

Each probe set was composed of six 1,2 to 1,5 kb-long PCR products, which were labeled using the FISH Tag DNA Multicolor kit (Alexa Fluor 488 dye for the *twist* promoter and Alexa Fluor 555 dye for the ectopic insertion sites) (Life Technologies). Mesodermal cells were stained using an anti-Twist antibody (anti-Rabbit polyclonal antibody generated by the Ghavi-Helm lab with assistance from the Protein Sciences Facility of the Lyon SFR Biosciences). Embryos were mounted in ProLong Gold antifade reagent with DAPI (Life Technologies) and imaged on a Leica SP8 confocal microscope using a 40x glycerol objective. For each embryo, several Z-stacks were acquired (section thickness of 0.361 μm) and processed using the Lightning Deconvolution software (Leica). A minimum of 500 nuclei from 3 to 4 independent embryos were analyzed and the relative distances between FISH signals measured using the Imaris software (Bitplane). A non-parametric two-sample Kolmogorov–Smirnov test was used to verify if the distance distributions were significantly different between samples. Two probes were considered co-localized when the distance between the centers of FISH signal was below 0.25 μm.

### Immunostaining

Immunostaining was performed as previously described^70^. The following primary antibodies were used: rabbit anti-Twist (1:200, generated by the Ghavi-Helm lab with assistance from the Protein Sciences Facility of the Lyon SFR Biosciences) and rat anti-TM1 (1:200, Developmental Studies Hybridoma Bank). Secondary antibodies were conjugated with Alexa 488 and Alexa 555 (Invitrogen). Confocal images were acquired using a Leica SP8 confocal microscope and processed using the Leica Application Suite X (LAS X) 3D Visualization and Adobe Photoshop CS6 software.

### smiFISH

smiFISH was performed as previously described^71^. Briefly, fixed embryos were incubated overnight at 37°C in hybridization buffer containing 320 nM of smiFISH probes. Embryos were washed and to immunostained with the appropriate antibody. Probes against *GFP* and *twist* were designed as previously described^71^. The X FLAP sequence was 5′ and 3′ end-labeled with Quasar 570. The embryos were mounted in ProLong Diamond antifade reagent with DAPI (Life Technologies) and imaged on a Leica SP8 confocal microscope using a 20x objective. The images were processed using the Adobe Photoshop CS6 software.

### Viability tests

Freshly hatched adult flies of the appropriate genotype were placed in embryo collection vials with standard apple cap plates and acclimatized at 25°C for 2 days prior to the experiments. After 3 pre-lays of 1 hour, eggs were collected for 2 hours at 25°C. Two hundred embryos were transferred on two new plates. Larvae and embryos were counted 24 hours later. Two independent experiments were performed for each genotype.

## DATA AVAILABILITY

All raw data were submitted to ArrayExpress (https://www.ebi.ac.uk/arrayexpress/browse.html) under accession numbers: E-MTAB-12153 (4C-seq), and E-MTAB-12146 (Micro-C).

## ACKNOWLEDGMENTS

We are very grateful to Mounia Lagha, Samir Merabet, Julie Soutourina, Michalis Averof, and François Leulier for critically reading the manuscript and to Frank Schnorrer for valuable input during the course of the study. We thank all members of the Ghavi-Helm lab for discussions and comments on the manuscript. This work was technically supported by the IGFL sequencing facility (PSI), the IGFL microscopy facility, the PLATIM imaging facility and Protein Sciences Facility of the Lyon SFR Biosciences (UAR3444/US8). This work was financially supported by an FRM starting grant (AJE20161236686) and ERC starting grant Enhancer3D (759708) to Y.G-H., an FRM doctoral fellowship (FDT202012010607) to A.G, and an internship fellowship from the UFR Biosciences Université Claude Bernard Lyon 1 to D.B.

## AUTHOR INFORMATION

### Contributions

Y.G-H. conceived and supervised the study. A.G., P.B.P., S.V., and Y.G-H. designed experiments. A.G. and S.V. generated all the transgenic lines with help from D.L. and H.T to perform microinjections. A.G., S.V., and Y.G-H. performed 4C-seq experiments. Y.G-H analyzed 4C-seq experiments. D.B performed and analyzed Micro-C experiments. P.B.P. performed and analyzed all imaging experiments. P.B.P. and S.V. performed viability tests. All of the authors discussed the results and implications and commented on the manuscript at all stages. Y.G-H. and A.G. acquired funding.

## ETHICS INTERESTS

### Competing interests

The authors declare no competing financial interests.

**Extended Data Figure 1:**
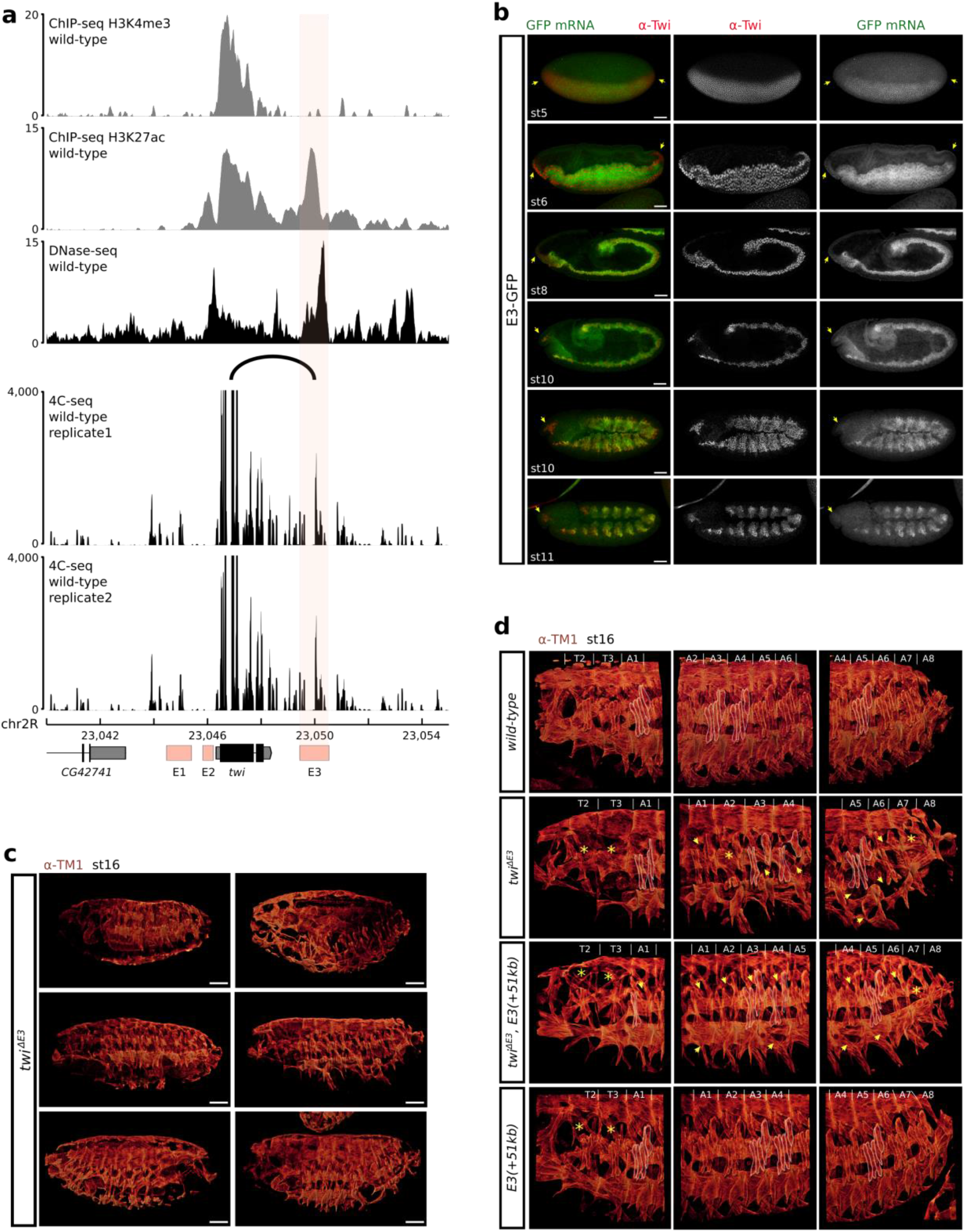
Characterization of the *twist* E3 enhancer. **a**. Top to bottom: ChIP-seq signal for histone modifications H3K4me3 and H3K27ac in sorted mesodermal cells at 6 to 8 hours after egg-lay^72^. DNase-seq signal in wild-type embryos at stage 11^42^. 4C-seq interaction maps at the *twist* locus in wild-type embryos at 5 to 8 hours after egg-lay. The observed interaction between the *twist* promoter and the E3 enhancer is highlighted by an arc. Two biological replicates are shown. **b**. Immunostaining with the α-Twist antibody (red) and expression (smiFISH) driven by its E3 enhancer (*GFP*, green) at stage 5, 6, 8, 10 (two different focal planes), and 11. The yellow arrows point to regions where E3 does not fully recapitulate the expression of *twist* (pole of the embryo during early stages and head region at later stages). Scale bars 50 μm. **c**. Immunostaining with the α-TM1 antibody at stage 16 (red, middle) in six different embryos of the *twi*^*ΔE3*^ line. Scale bars 50 μm. d. Immunostaining with the α-TM1 antibody at stage 16 (red, middle) in wild-type, *twi*^*ΔE3*^, *twi*^*ΔE3*^, *E3(+51kb)*, and *E3(+51kb)* embryos. The location of specific embryonic body muscles is highlighted in white and the location of missing muscles in the mutants in yellow (yellow asterisk: muscles absent in the whole segment, yellow arrows: absence of specific muscles).

**Extended Data Figure 2:**
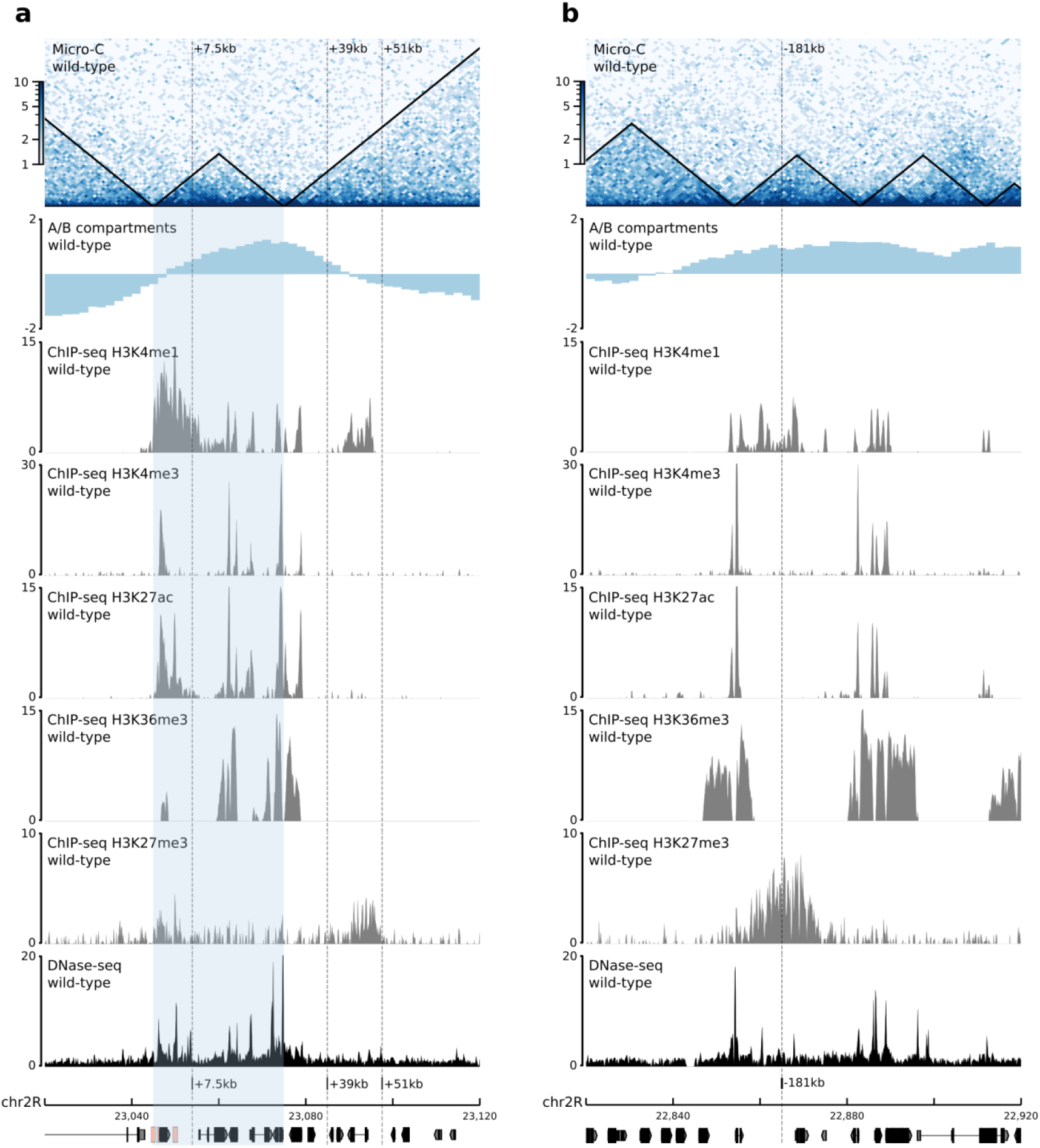
Chromatin organization surrounding the +7.5kb, +39kb, +51kb (a), and -181kb (b) insertion sites. **a. b**. Top to bottom: Normalized Micro-C contact map at 1000 bp resolution in wild-type embryos at 5 to 8 hours after egg-lay (two biological replicates merged). A/B compartments in wild-type embryos at stage 5-8^43^. ChIP-seq signal for histone modifications H3K4me1, H3K4me3, H3K27ac, H3K36me3, and H3K27me3 in sorted mesodermal cells at 6 to 8 hours after egg-lay^72^. DNase-seq signal in wild-type embryos at stage 11^42^. The TAD containing *twist* and the E3 enhancer is highlighted in light blue.

**Extended Data Figure 3:**
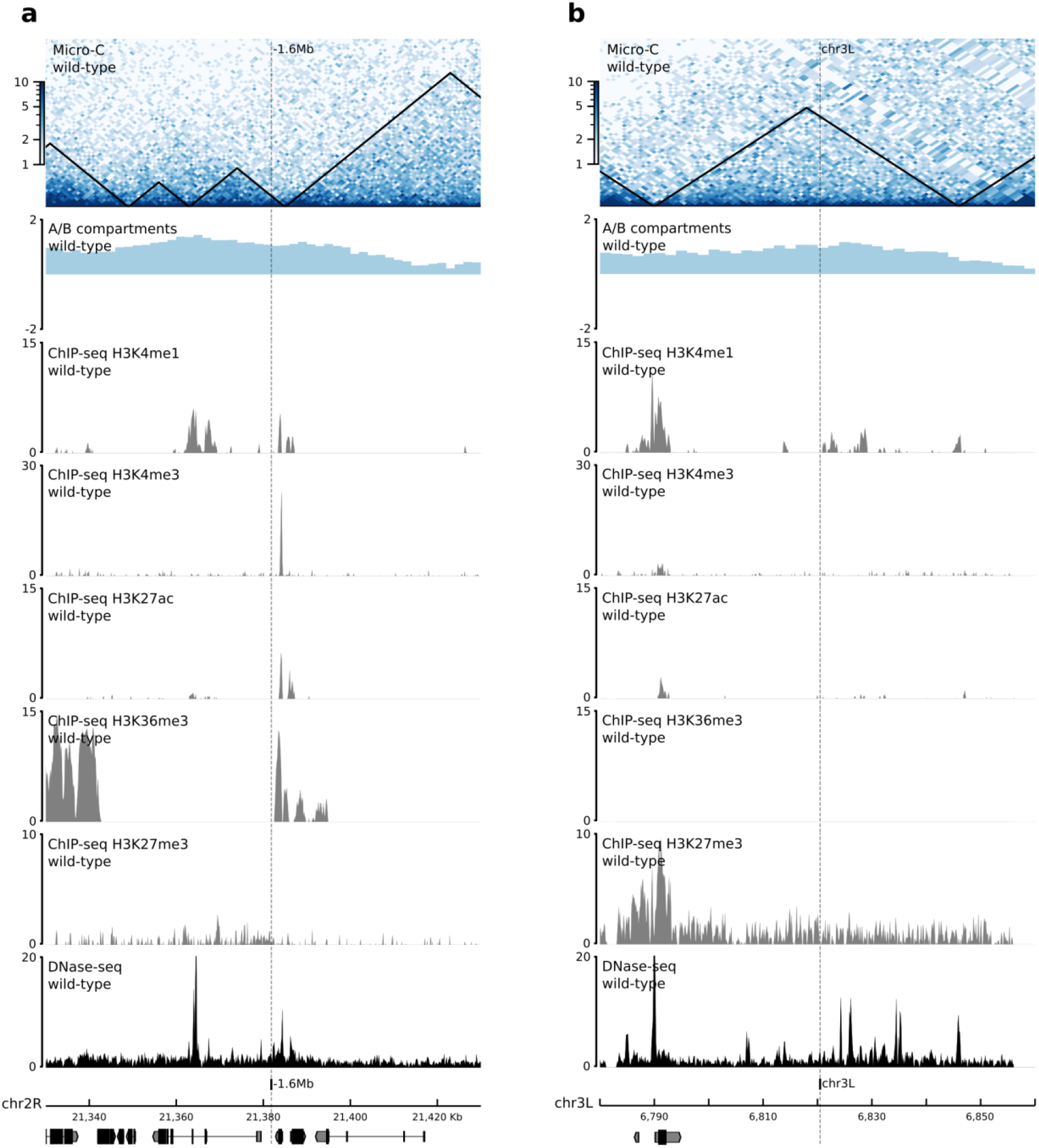
Chromatin organization surrounding the -1.6Mb (a), and chr3L (b) insertion sites. **a. b**. Top to bottom: Normalized Micro-C contact map at 1000 bp resolution in wild-type embryos at 5 to 8 hours after egg-lay (two biological replicates merged). A/B compartments in wild-type embryos at stage 5-8^43^. ChIP-seq signal for histone modifications H3K4me1, H3K4me3, H3K27ac, H3K36me3, and H3K27me3 in sorted mesodermal cells at 6 to 8 hours after egg-lay^72^. DNase-seq signal in wild-type embryos at stage 11^42^.

**Extended Data Figure 4:**
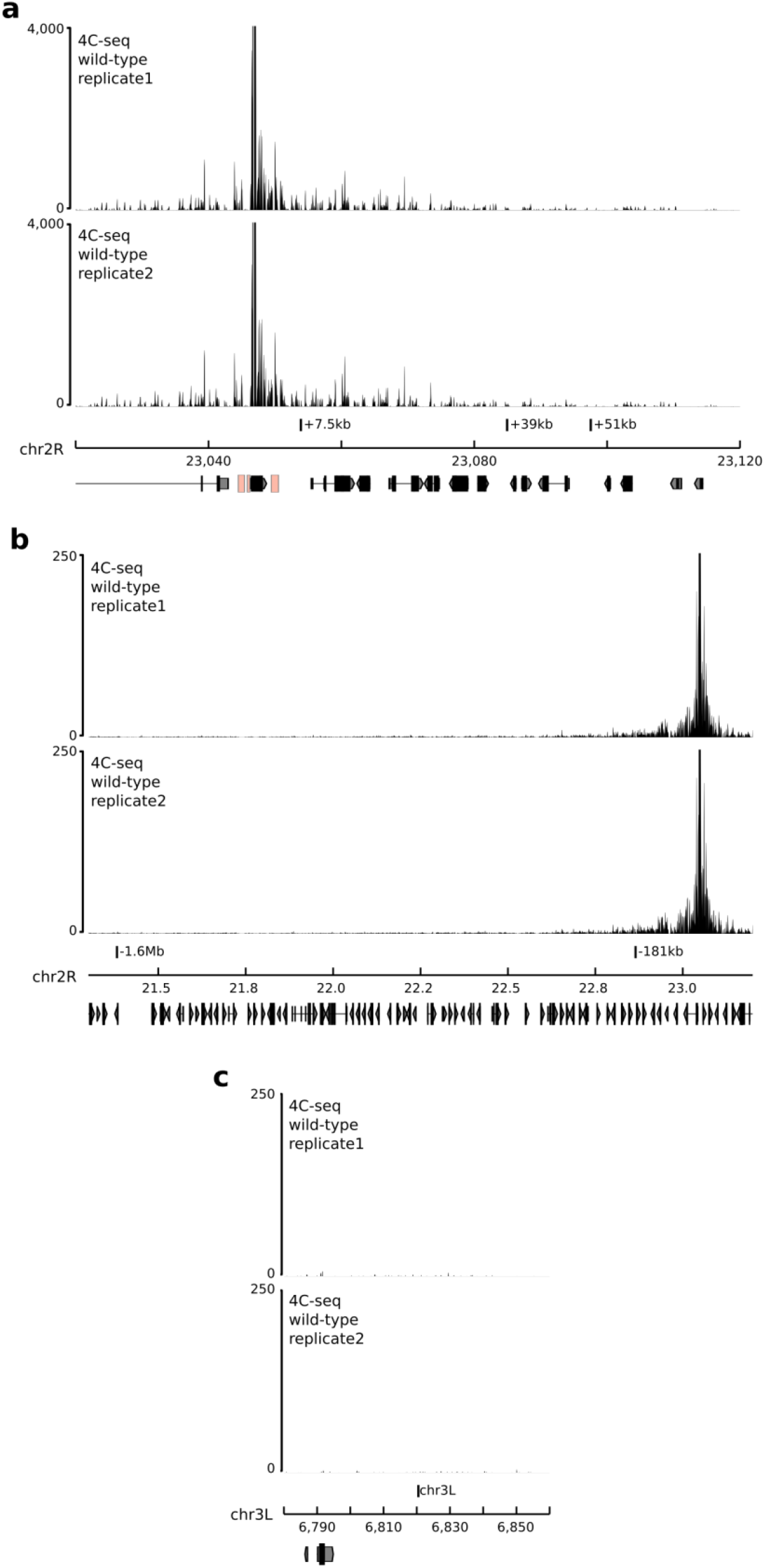
The selected insert sites do not interact with the *twist* locus in wild-type embryos. **a. b. c**. 4C-seq interaction maps around the +7.5kb, +39kb, +51kb (a), -181kb, -1.6MB (b), and chr3L (c) insert sites in wild-type embryos at 5 to 8 hours after egg-lay. Two biological replicates are shown.

**Extended Data Figure 5:**
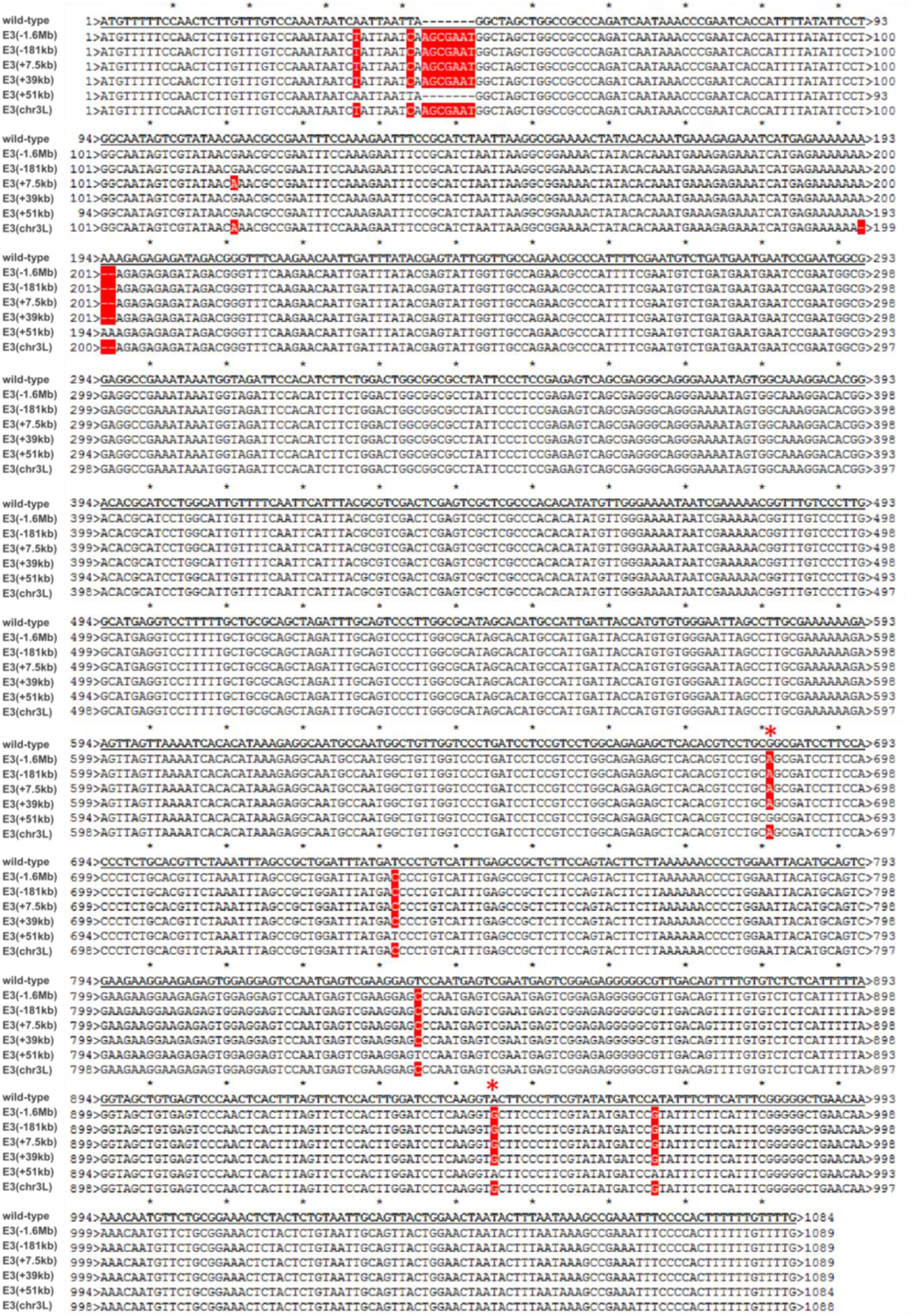
Alignment of the wild-type E3 enhancer sequence with the ectopic E3 enhancers from the different transgenic lines. Sequence alignment of the E3 enhancer inserted at the ectopic sites (wild-type), and of the endogenous E3 sequence present in the different transgenic lines. Note that in the case of line *E3(+51kb)*, the endogenous sequence is that of the wild-type sequence, while the ectopic sequence is that of the E3(−1.6Mb) sequence. Mismatches are highlighted in red. A red asterisk indicated the location of the two variants used to calculate the percentage of ectopic E3 reads (Methods).

**Extended Data Figure 6:**
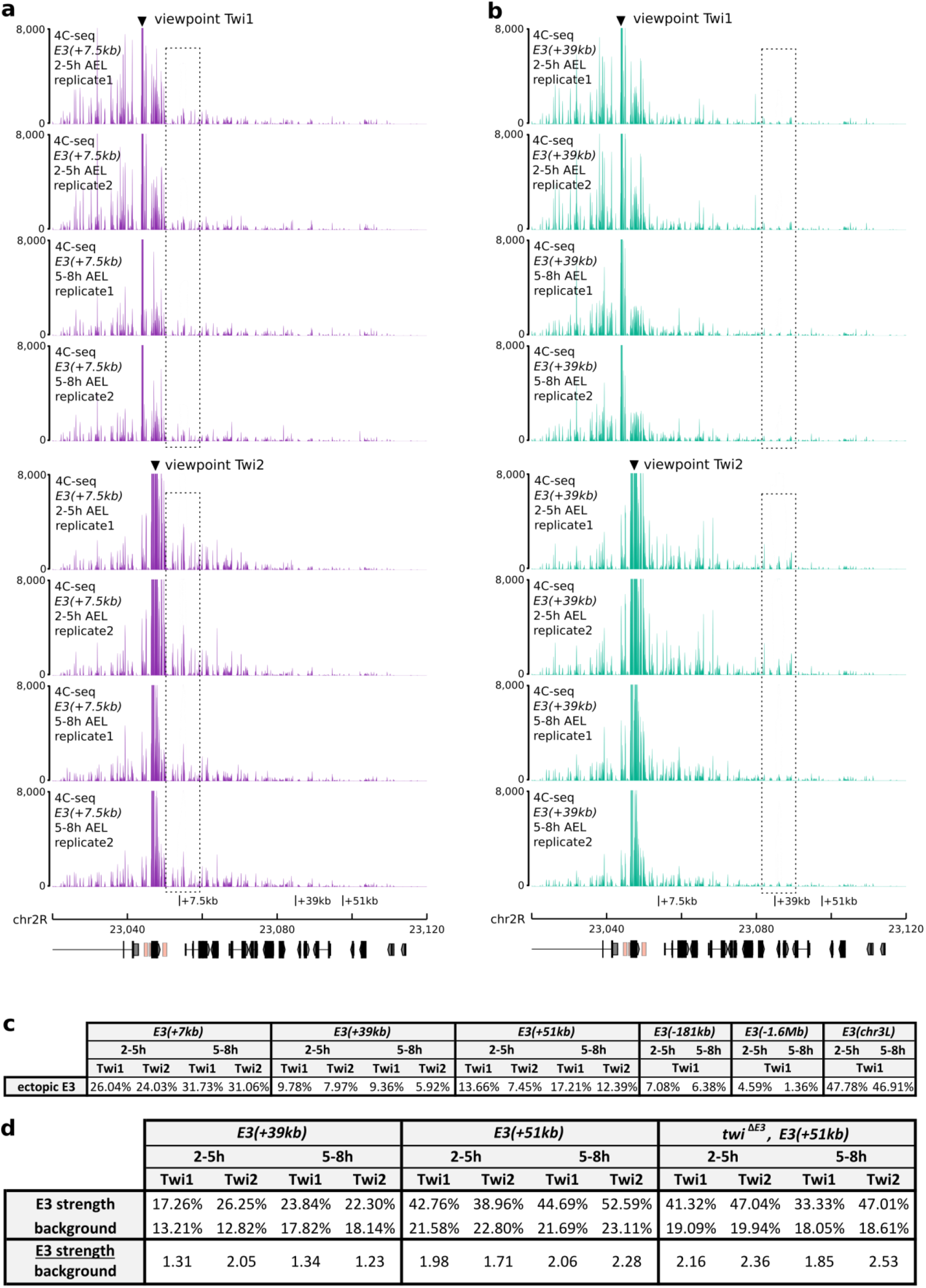
Analysis of the long-range interactions between the *twist* promoter and E3(+7.5kb) and E3(+39kb) ectopic enhancers. **a. b**. 4C-seq interaction maps around the +7.5kb, +39kb, and +51kb insert sites in *E3(+7*.*5kb)* (**a**.) and *E3(+39kb)* (**b**.) embryos at 2 to 5 hours and 5 to 8 hours after egg-lay (AEL). Two biological replicates and two different viewpoints are shown. **c**. Percentage of ectopic E3 reads in 4C-seq datasets from *E3(+7*.*5kb), E3(+39kb), E3(+51kb), E3(−181kb), E3(−1*.*6Mb)*, and *E3(chr3L)* embryos at 2 to 5 hours and 5 to 8 hours after egg-lay for each viewpoint. **d**. Percentage of reads mapping on the 2-kb ectopic E3 over a 10-kb window (E3 strength), percentage of reads mapping on an adjacent control region (background), and the ratio between there percentages in 4C-seq datasets from *E3(+39kb), E3(+51kb)*, and *twi*^*ΔE3*^, *E3(+51kb)* embryos at 2 to 5 hours and 5 to 8 hours after egg-lay for each viewpoint.

**Extended Data Figure 7:**
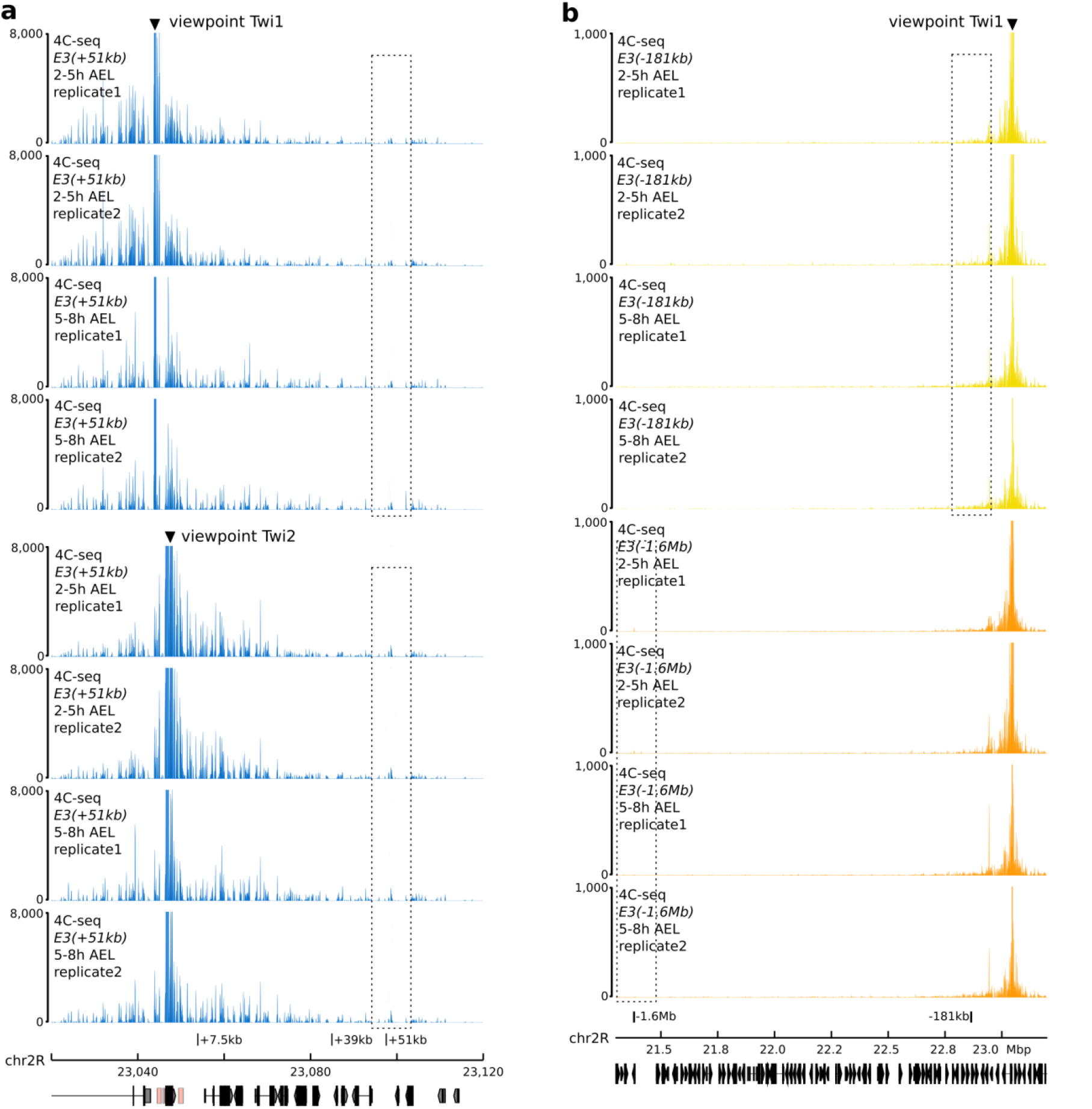
Analysis of the long-range interactions between the *twist* promoter and E3(+51kb) and E3(−181kb) and E3(−1.6Mb) ectopic enhancers. **a**. 4C-seq interaction maps around the +7.5kb, +39kb, and +51kb insert sites in *E3(+51kb)* embryos at 2 to 5 hours and 5 to 8 hours after egg-lay (AEL). Two biological replicates and two different viewpoints are shown. **b**. 4C-seq interaction maps around in *E3(−181kb)* (top, yellow) *E3(−1*.*6Mb)* (bottom, orange) embryos at 2 to 5 hours and 5 to 8 hours after egg-lay. Two biological replicates are shown.

**Extended Data Figure 8:**
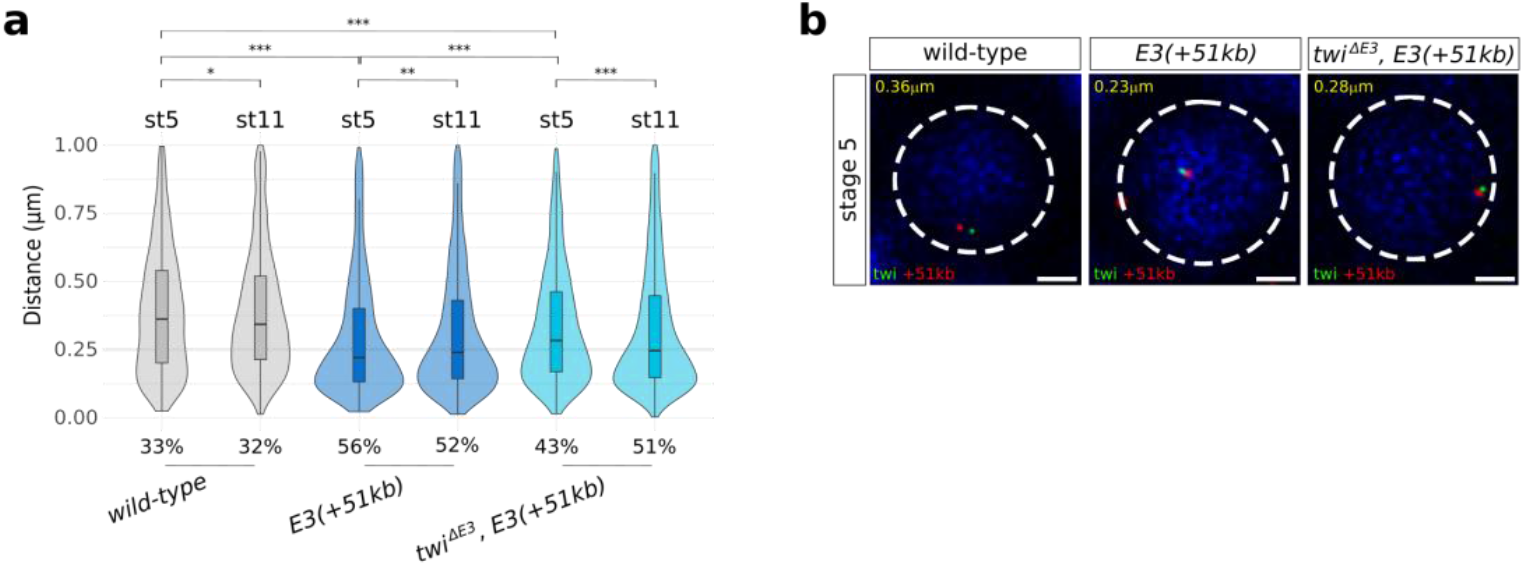
3D DNA FISH a stage 5. **a**. Violin plots representing 3D DNA FISH distances measured in mesodermal nuclei between a probe located next to the *twist* promoter and a probe located next to the +51kb insert site in wild-type (grey), *E3(+51kb)* (dark blue), and *twi*^*ΔE3*^, *E3(+51kb)* (light blue) embryos at stage 5 and 11. A non-parametric two-sample Kolmogorov–Smirnov test was used to assess the significant difference between DNA FISH distance distributions (wild-type stage 5 versus wild-type stage 11: p = 0.045, *E3(+51kb)* stage 5 versus *E3(+51kb)* stage 11: p = 0.011, *twi*^*ΔE3*^, *E3(+51kb)* stage 5 versus *twi*^*ΔE3*^, *E3(+51kb)* stage 11: p = 4.61e^-7^, wild-type stage 5 versus *E3(+51kb)* stage 5: p = 2.2e^-16^, wild-type stage 5 versus *twi*^*ΔE3*^, *E3(+51kb)* stage 5: p = 4.62e^-10^, *E3(+51kb)* stage 5 versus *twi*^*ΔE3*^, *E3(+51kb)* stage 5: p = 4.06e^-14^). The percentage of colocalization (defined as the percentage of probe pairs with a distance < 0.25 μm; Methods) is indicated for each condition. **b**. Representative pictures of DNA FISH nuclei from a. Scale bars 1 μm.

**Extended Data Figure 9:**
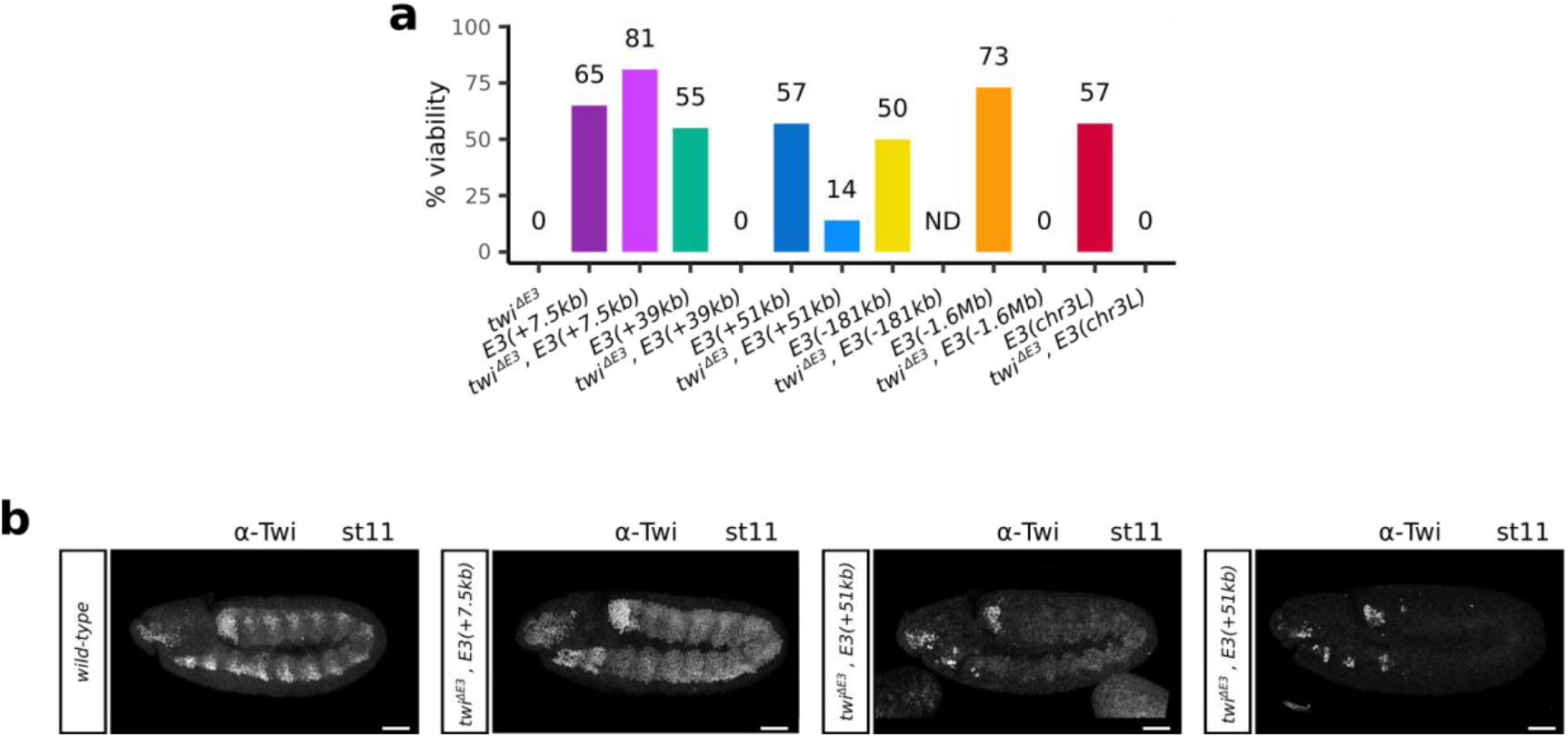
Characterization of transgenic lines. **a**. Bar plot representing the percentage of viable embryos in the different fly lines under study. For each condition, at least two independent experiments were performed, with at least 50 embryos each. ND: Not Determined. **b**. Immunostaining with the α-Twist antibody at stage 11 in wild-type, *twi*^*ΔE3*^, *E3(+7*.*5kb)*, and *twi*^*ΔE3*^, *E3(+51kb)* embryos. Scale bars 50 μm.

**Extended Data Figure 10:**
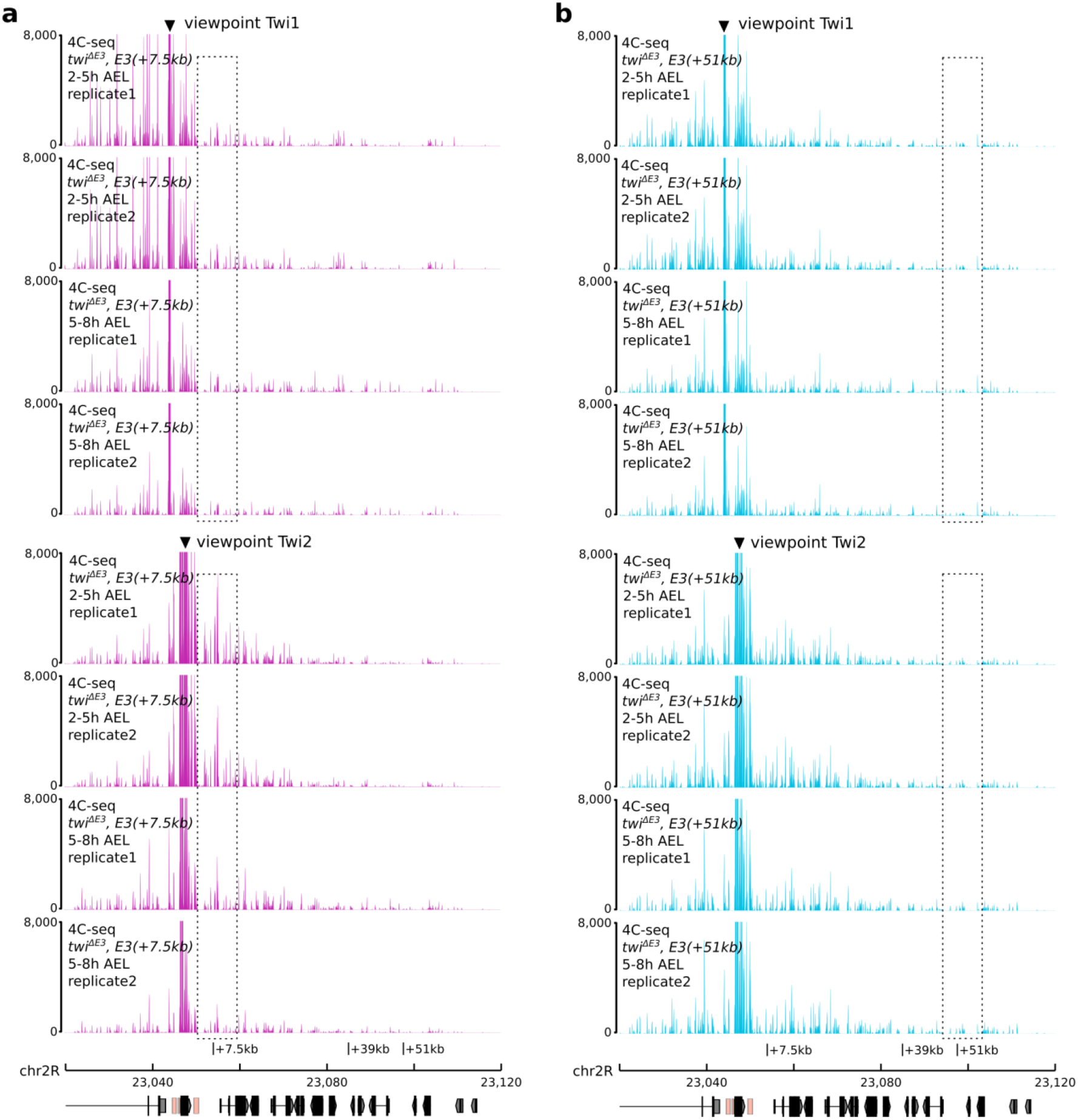
Analysis of the long-range interactions in *twi*^*ΔE3*^, *E3(+7*.*5kb)* and *twi*^*ΔE3*^, *E3(+51kb)* embryos. **a. b**. 4C-seq interaction maps around the +7.5kb, +39kb, and +51kb insert sites in *twi*^*ΔE3*^, *E3(+7*.*5kb)* (**a**.) and *twi*^*ΔE3*^, *E3(+51kb)* (**b**.) embryos at 2 to 5 hours and 5 to 8 hours after egg-lay (AEL). Two biological replicates and two different viewpoints are shown.

**Extended Data Figure 11:**
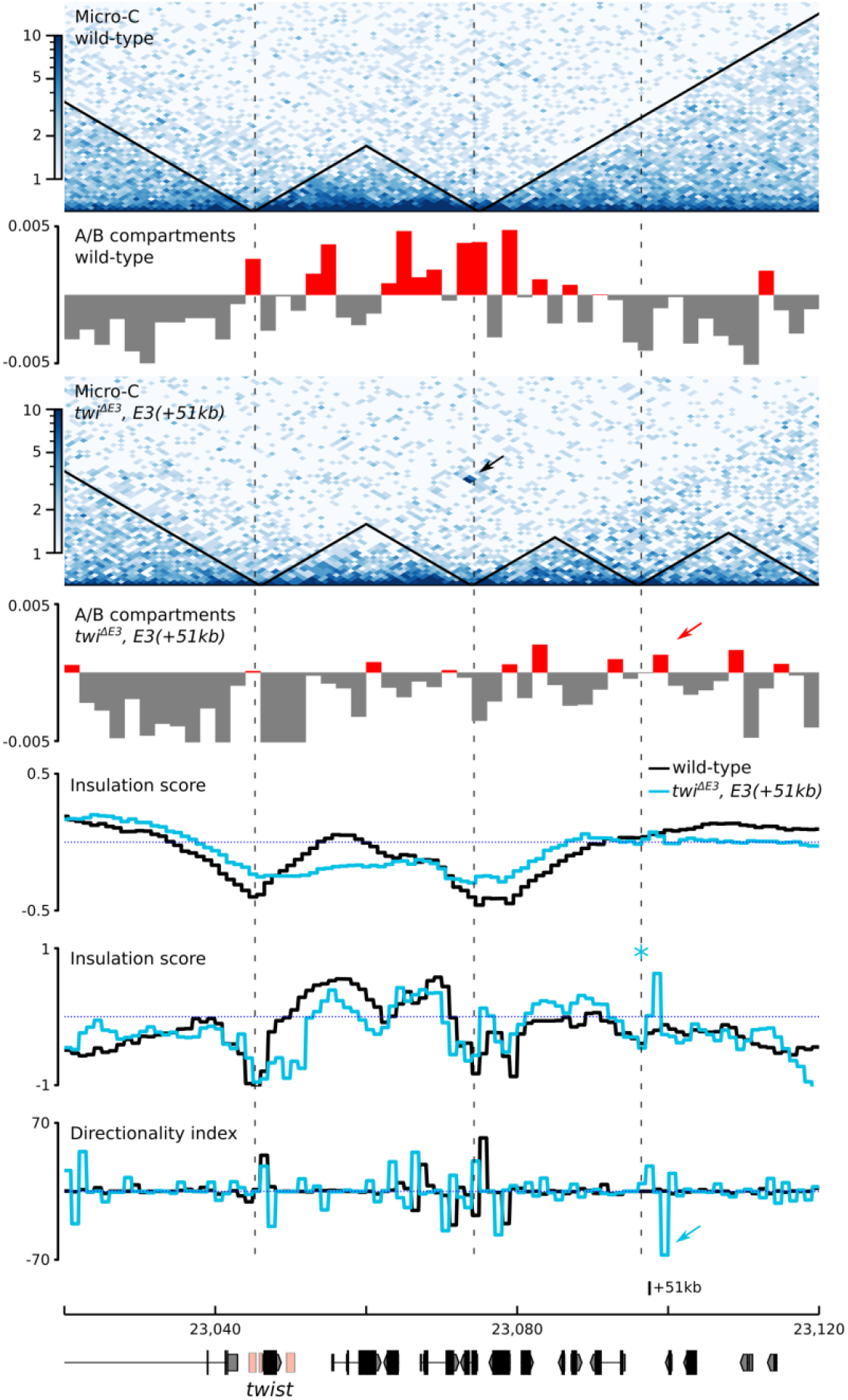
Chromatin organization surrounding the +51kb insertion site in wild type and *twi*^*ΔE3*^, *E3(+51kb)* embryos. Top to bottom: Normalized Micro-C contact map at 1000 bp resolution in wild-type embryos at 5 to 8 hours after egg-lay (two biological replicates merged). A (red) / B (grey) compartments in wild-type embryos at 5 to 8 hours after egg-lay. Normalized Micro-C contact map at 1000 bp resolution in *twi*^*ΔE3*^, *E3(+51kb)* embryos at 5 to 8 hours after egg-lay (two biological replicates merged). A (red) / B (grey) compartments in *twi*^*ΔE3*^, *E3(+51kb)* embryos at 5 to 8 hours after egg-lay. The interaction between the *twist* locus and the +51kb insertion site is indicated by a black arrow. The location of TAD boundaries is indicated by vertical dotted lines. The shift in compartment in *twi*^*ΔE3*^, *E3(+51kb)* embryos is indicated by a red arrow. Insulation score computed using HiCexplorer in wild-type (black) and *twi*^*ΔE3*^, *E3(+51kb)* (light blue) embryos. Insulation score computed using FAN-C in wild-type (black) and *twi*^*ΔE3*^, *E3(+51kb)* (light blue) embryos. Note the overall similar insulation landscape, specifically the maintenance of the TAD boundary between the *twist* locus and the +51 insertion site, and the appearance of a new boundary around the +51 insertion site in *twi*^*ΔE3*^, *E3(+51kb)* embryos (light blue asterisk). Directionality index computed using FAN-C in wild-type (black) and *twi*^*ΔE3*^, *E3(+51kb)* (light blue) embryos. Note the appearance of a strong negative shift in the directionality index around the +51 insertion site in *twi*^*ΔE3*^, *E3(+51kb)* embryos (light blue arrow), indicative of biased interaction towards the left.

**Extended Data Figure 12:**
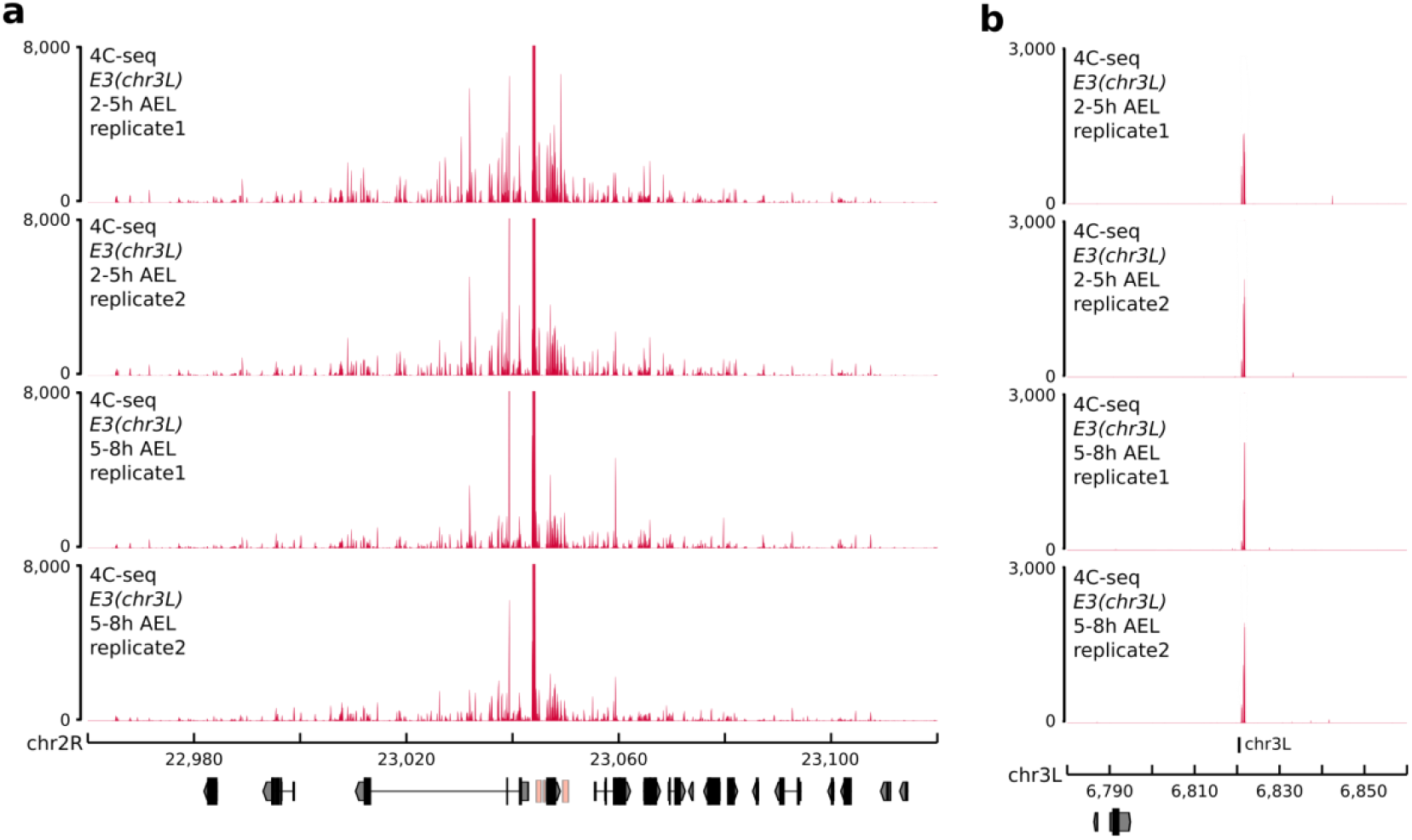
Analysis of the long-range interactions between the *twist* promoter and E3(chr3L) ectopic enhancer. **a**. 4C-seq interaction maps around the *twist* locus in *E3(chr3L)* embryos at 2 to 5 hours and 5 to 8 hours after egg-lay (AEL). Two biological replicates are shown. **b**. 4C-seq interaction maps around chr3L insert site in *E3(chr3L)* embryos at 2 to 5 hours and 5 to 8 hours after egg-lay. Two biological replicates are shown.

